# Efficient population coding depends on stimulus convergence and source of noise

**DOI:** 10.1101/2020.06.15.151795

**Authors:** Kai Röth, Shuai Shao, Julijana Gjorgjieva

## Abstract

Sensory organs transmit information to downstream brain circuits using a neural code comprised of spikes from multiple neurons. According to the prominent efficient coding framework, the properties of sensory populations have evolved to encode maximum information about stimuli given biophysical constraints. How information coding depends on the way sensory signals from multiple channels converge downstream is still unknown, especially in the presence of noise which corrupts the signal at different points along the pathway. Here, we calculated the optimal information transfer of a population of nonlinear neurons under two scenarios. First, a lumped-coding channel where the information from different inputs converges to a single channel, thus reducing the number of neurons. Second, an independent-coding channel when different inputs contribute independent information without convergence. In each case, we investigated information loss when the sensory signal was corrupted by two sources of noise. We determined critical noise levels at which the optimal number of distinct thresholds of individual neurons in the population changes. Comparing our system to classical physical systems, these changes correspond to first- or second-order phase transitions for the lumped- or the independent-coding channel, respectively. We relate our theoretical predictions to coding in a population of auditory nerve fibers recorded experimentally, and find signatures of efficient coding. Our results yield important insights into the diverse coding strategies used by neural populations to optimally integrate sensory stimuli in the presence of distinct sources of noise.

## Introduction

Neurons in sensory organs encode information about the environment and transmit it to downstream circuits in the brain. In many sensory systems, the sensory signal is not coded merely by individual neurons but rather by the joint activity of populations of neurons, which likely coordinate their responses to represent the stimulus as efficiently as possible. One signature of this efficient parallel coding might be the remarkably diverse response properties exhibited by many first-order sensory neurons. In the visual pathway, for example, the first-order sensory neurons are the retinal ganglion cells (RGCs) which send information to the thalamus through the optic nerve. There exist around thirty different RGC types which encode different visual features as characterized by the cells’ spatiotemporal receptive fields and nonlinear computations [1–3]. Yet, there are also RGC types which in parallel encode a single stimulus feature differing in their firing thresholds [4–6], and hence provide parallel information streams. Another example is the first synapse level of the auditory pathway, where each inner hair cell transmits information about sound intensity to approximately ten to thirty different auditory nerve fibers (ANFs) [7]. ANFs differ in several aspects of their responses, including spontaneous rates and firing thresholds [8]. However, each fiber receives exclusive input from only a single inner hair cell. As in the retina, this results in a highly parallelized stream of sensory information. Similarly, this parallel encoding of a single stimulus feature with a population of neurons with different thresholds has been shown in olfactory receptor neurons [9], in mammalian touch receptors [10], and electro receptors of electric fish [11].

We asked whether the diverse response properties of a population of neurons encoding a single stimulus feature are a consequence of the evolutionary pressure of the sensory system to efficiently encode sensory stimuli. A powerful theoretical framework to address this question is the efficient coding. This framework postulates that during evolution sensory systems have optimized information encoding given biophysical and metabolic constraints. Predictions from efficient coding are consistent with many properties of primary sensory neurons, including center-surround receptive fields [12] and a split into ON and OFF pathways in the retina [13,14], as well as the input-output functions of neurons [15] and sensory adaptation to changing stimulus statistics in the insect retina [16]. Applying the efficient coding framework requires determining a set of constraints that are relevant for the sensory system in question. Rather than investigating efficient coding in a specific sensory system, we sought to derive a general theoretical framework that applies to multiple sensory systems focusing on two questions: first, how the source and size of noise affects the accuracy of information coding, and second, how downstream signal convergence influences the optimality of information transfer.

Noise is a ubiquitous phenomenon in biological information processing and corrupts signal transmission at different processing stages (reviewed in [17]). The size [18–24] and source [25,26] of noise can have distinct effects on signal encoding. For instance, previous studies have shown that neural populations adopt a strategy of independent coding [5,14,25,27–30] or decorrelation [12,31–37] in conditions of low noise, and a strategy of redundant coding in the presence of high noise [5,12,14, 25, 27–35, 37, 38]. For populations of neurons, redundant coding can be interpreted as multiple neurons in the population, which acquire the same response thresholds to average out uncertainties in stimulus representation due to noise. When noise is negligible, the individual thresholds are expected to be distinct from each other, which is in agreement with experimental data [5, 28, 30]. Therefore, the source and size of noise have nontrivial influences on the encoding of sensory information.

Besides the source of noise, a second factor when maximizing information between stimulus and response is how the stimulus converges downstream after it is encoded by the neural population. For instance, it might be advantageous to compress sensory information before reaching downstream circuits due to axonal transmission limitations and the metabolic cost of firing of multiple neurons. Previous studies have assumed a framework in which the spiking output of the neurons converges, or is lumped, into one single output variable [29,30]. In contrast, other works have assumed a framework without signal convergence, i.e. where the signal is encoded by the independent spiking output of each neuron in the population [5, 14, 25, 39]. Therefore, signal convergence also fundamentally influences optimal population coding.

Here, we investigate efficient stimulus coding in populations of more than two neurons as a function of the source and size of noise, and the type of stimulus convergence, and generate novel predictions about how these two aspects affect the optimal coding strategies of the neural populations. In particular, we maximize Shannon’s mutual information between a one-dimensional stimulus (a single stimulus feature) and the population’s response. Typically, the efficient coding framework has been applied to populations of linear neurons, where the contribution of the noise entropy term to the mutual information has been ignored, resulting in degeneracies in the optimal solutions [12,13,37,40]. We consider a nonlinear version of efficient coding that includes two types of noise: additive input noise which corrupts the stimulus before it enters a neuron’s nonlinearity and output noise implemented as spike generating noise which affects the output of the nonlinearity. We found that the exact implementation and source of noise can have fundamental implications for the conclusions arising from efficient coding.

For biologically realistic intermediate sources of noise, the particular downstream convergence of the sensory signal, either through lumping or independence, determines the number of distinct population thresholds. In agreement with previous studies [5,14,25,27–30], we found that for low noise levels the optimal population thresholds are all distinct, while for high noise levels all thresholds become identical. However, unlike other studies we found surprising transitions from all thresholds being distinct at low noise to all thresholds being equal at high noise, which happen through a set of bifurcations at critical noise levels. These bifurcations resemble first- and second-order phase transitions in the case of the lumped and independent output variables, respectively. We related these phase transitions to the curvature of the information landscape, giving us insights into the optimal coding solutions and their relationship to well-understood critical systems in physics. We also compared our theoretical predictions to optimal coding of experimentally recorded auditory nerve fibers where we found signatures of optimality.

## Results

### Theoretical framework

We studied a population of spiking neurons encoding a sensory stimulus under different noise scenarios. A population of *N* neurons encodes a static, one-dimensional stimulus *s* drawn from a stimulus distribution *P*(*s*) through the spike counts 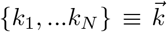 emitted in a coding time window Δ*T* (Fig. 1A). We considered different stimulus distributions, parametrized by the generalized normal distribution [41], but here we primarily discuss the case of a Gaussian stimulus distribution (see Methods). The mapping from the stimulus value *s* to the spike count vector 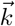 happens through a set of *N* nonlinear functions (tuning curves) {*ν*_1_(*s*),…*ν_N_*(*s*)}, where *ν_i_*(*s*) denotes the firing rate of the respective neuron *i*. Such a mapping can be implemented by a variety of sensory systems, for instance, the retina which processes various visual stimulus attributes, such as light intensity or contrast [42], the olfactory receptor neurons which process a range of concentrations of a single odor [9, 39, 43, 44], or the auditory nerve fibers (ANFs) which transmit information about sound pressure levels [45]. Here, we only focus on optimizing one stage of this transformation, namely the nonlinearity which takes a filtered stimulus *s* as input and converts it into action potentials 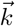. For example, if we assume that our sensory system of interest is the population of ANFs, which are highly nonlinear processing units [46], then *s* represents the stimulus value following preprocessing by the cochlea and the inner hair cells.

**Figure 1.**
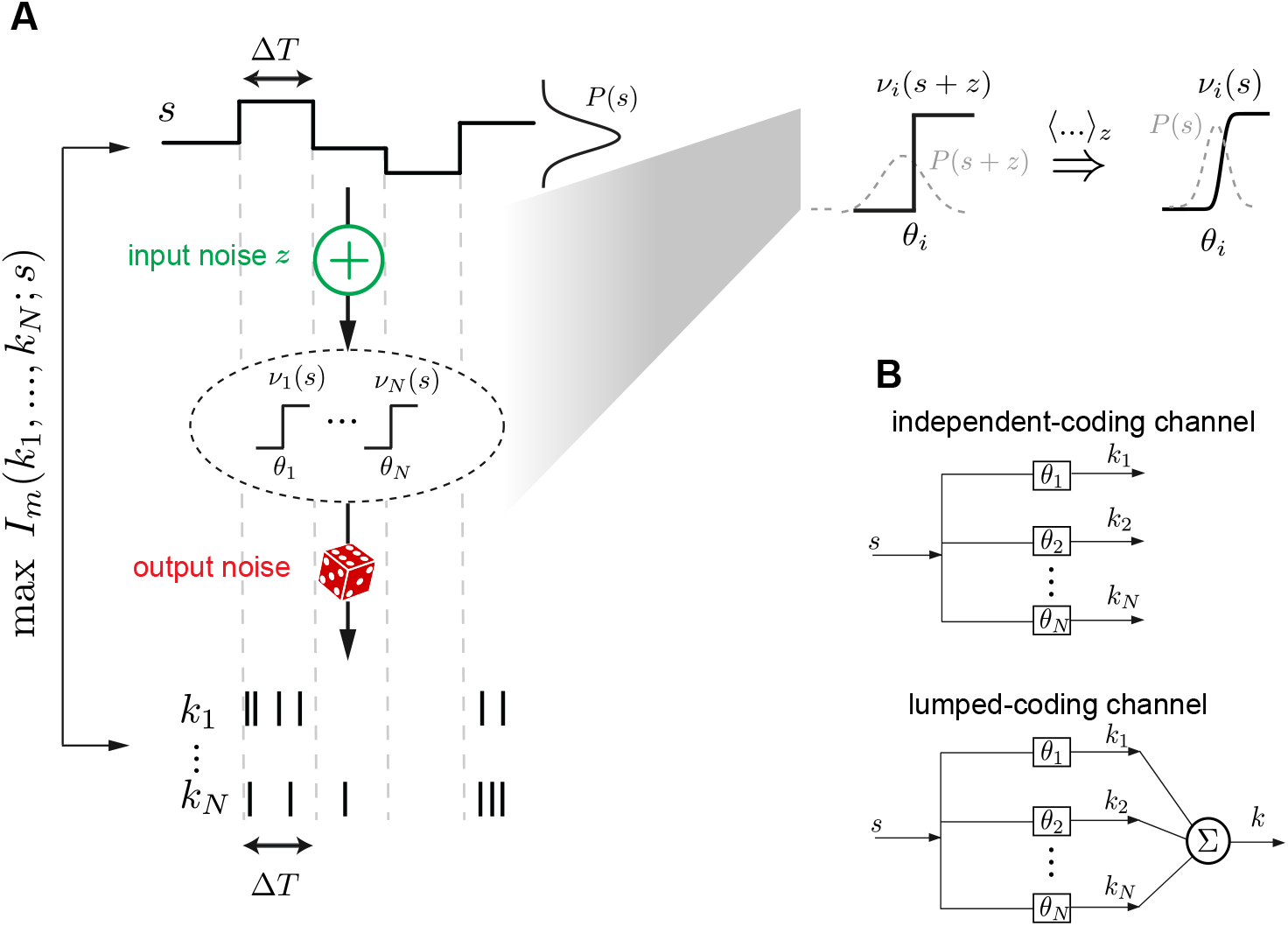
Stimulus encoding with a population of neurons in the presence of input and output noise. **A.** Framework: A static stimulus *s* (top) is encoded by a population of spike counts {*k*_1_,…*k_N_*} (bottom) in a coding time window Δ*T*. The stimulus is first corrupted by additive input noise *z* and then processed by a population of *N* binary nonlinearities {*ν*_1_,…*ν_N_*}. Stochastic spike generation based on Poisson output noise corrupts the signal again. Thresholds {*θ*_1_,…,*θ_N_*} of the nonlinearities are optimized such that the mutual information *I_m_*(*k*_1_,…, *k_N_*; *s*) between stimulus and spike counts is maximized. Inset: Introducing additive input noise and a binary nonlinearity can be interpreted as having a sigmoidal nonlinearity after the input noise is averaged, 〈⋯〉_*z*_. Shallower nonlinearities result from higher input noise levels. **B.** Two different scenarios of information transmission: In the independent-coding channel each neuron contributes with its spike count to the coding of the stimulus, while in the lumped-coding channel all spike counts are added into one scalar output variable that codes for the stimulus.

We modeled the neurons’ tuning curves as binary, described by two firing rate levels {0, *ν*_max_} with an individual threshold *θ_i_* separating the stimuli into two firing rates. Thus, the input-output functions of each neuron can be represented by *ν_i_*(*x*) = *ν*_max_Θ(*θ_i_* − *x*), where Θ is the Heaviside function. This simplification is justified by the fact that many sensory neurons have been described with steep tuning curves that resemble binary neurons [16,27,36], and it makes the problem mathematically traceable. We derived the number and values of distinct thresholds in the population when the signal is corrupted by two sources of noise: input noise, which affects the signal before the nonlinearity, and output noise, which affects the neuronal outputs after the nonlinearity.

#### Input noise

Before being processed by the nonlinearity, the stimulus *s* is corrupted by additive noise *z* drawn from a distribution *P*(*z*). The size of input noise can be quantified by the ratio of its variance 〈*z*^2^〉 ≡ *σ*^2^ to the stimulus variance 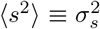. Without loss of generality, we set 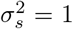 and thus *σ*^2^ alone stands for the size of input noise. The noise affects the stimulus independently for each nonlinearity; we did not consider correlated noise since previous work has shown that the case of correlated noise can be reduced to independent noise with lower *σ*^2^ [25]. Similarly to the stimulus distribution, we primarily examined the case with the noise drawn from a Gaussian distribution, *z* ~ *N*(0, *σ*^2^), but we also considered other distributions (see Methods). Since the input to the nonlinearities is *x* = *s* + *z*, the effective tuning curves, *ν_i_*(*s*), can be described to have sigmoidal shape (Fig. 1A, inset). A larger input noise size, determined by the variance of the noise *σ*^2^, corresponds to a shallower slope. In the remainder of the text, we use the standard deviation *σ* to refer to the size of input noise.

#### Output noise

Output noise was implemented by generating output spikes stochastically; here, each of the spike counts *k_i_* in a coding window Δ*T* was Poisson distributed. Large output noise corresponds to the case when the product of *ν*_max_ and Δ*T* is small; in this case the output of a given cell *i* is often *k_i_* = 0 making it more difficult to distinguish whether the underlying firing rate for that neuron is 0 and thus the stimulus is smaller than the threshold *θ_i_*, or whether the firing rate is *ν*_max_ and the stimulus greater than *θ_i_*. The output noise size can thus be quantified by the expected spike count for maximum firing rate, *R*: = *ν*_max_Δ*T*, where small *R* means high noise.

Within this framework, we maximized the mutual information between stimulus and output spike counts, and optimized the number and values of distinct thresholds, {*θ_i_*}, of the neuronal nonlinearities, while varying the size of input and output noise (see Methods). We chose the mutual information as the objective function to quantify the optimality of the encoding because it does not rely on any specific assumptions of how this information should be decoded, and presents an upper bound for any other efficiency measure [47]. In addition to two noise sources, we further distinguish between two different scenarios previously considered in the literature for how the sensory signal converges after being processed by the population of neurons (Fig. 1B): (1) an *independent-coding channel* where a vector of spike counts 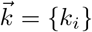 generates a population code of the stimulus where each spike count independently contributes to the total information [5,14,25], and (2) a *lumped-coding channel* where a scalar output variable 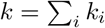, obtained by summing the individual spike counts *k_i_*, codes for the stimulus [29,30].

### The independent-coding channel transmits more information than the lumped-coding channel

To understand how stimulus convergence influences information transmission in larger neural populations of more than two neurons, we compared the mutual information and optimal thresholds between the lumped- and independent-coding channel scenarios in the presence of two noise sources.

To first gain intuition, we illustrate the case with vanishing input noise (*σ* = 0) and a population with two neurons with thresholds *θ*_1_ < *θ*_2_, which divide the entire stimulus distribution into three regions: Δ_1_: *s* < *θ*_1_, Δ_2_: *θ*_1_ ≤ *s* < *θ*_2_ and Δ_3_: *s* ≥ *θ*_2_ (Fig. 2, left). Here, we compute all possible spike counts and corresponding “estimation probabilities,” 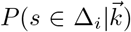, which describe the probability of the stimulus being in each of the three regions {Δ_*i*_}_*i*={1,2,3}_ for a given spike count 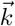 (Fig. 2). These vary as a function of output noise, and we consider three cases: high, intermediate, and negligible output noise. First, in the limit of vanishing output noise where *R* = *ν*_max_Δ*T* is very large, the information encoded by both channels is identical because both reach capacity and transmit log_2_(3) bits of information (Fig. 2A). In particular, whenever the stimulus is larger than the threshold of a given cell, that cell will on average fire R spikes. Since *R* → ∞, for that given cell the probability of having 0 spikes is infinitesimal. This unambiguously determines the stimulus region {Δ_*i*_} in which the stimulus occurs. Hence, the estimation probabilities all become either 0 or 1, leading to identical output entropy for both coding channels, and consequently identical mutual information with zero noise entropy.

**Figure 2.**
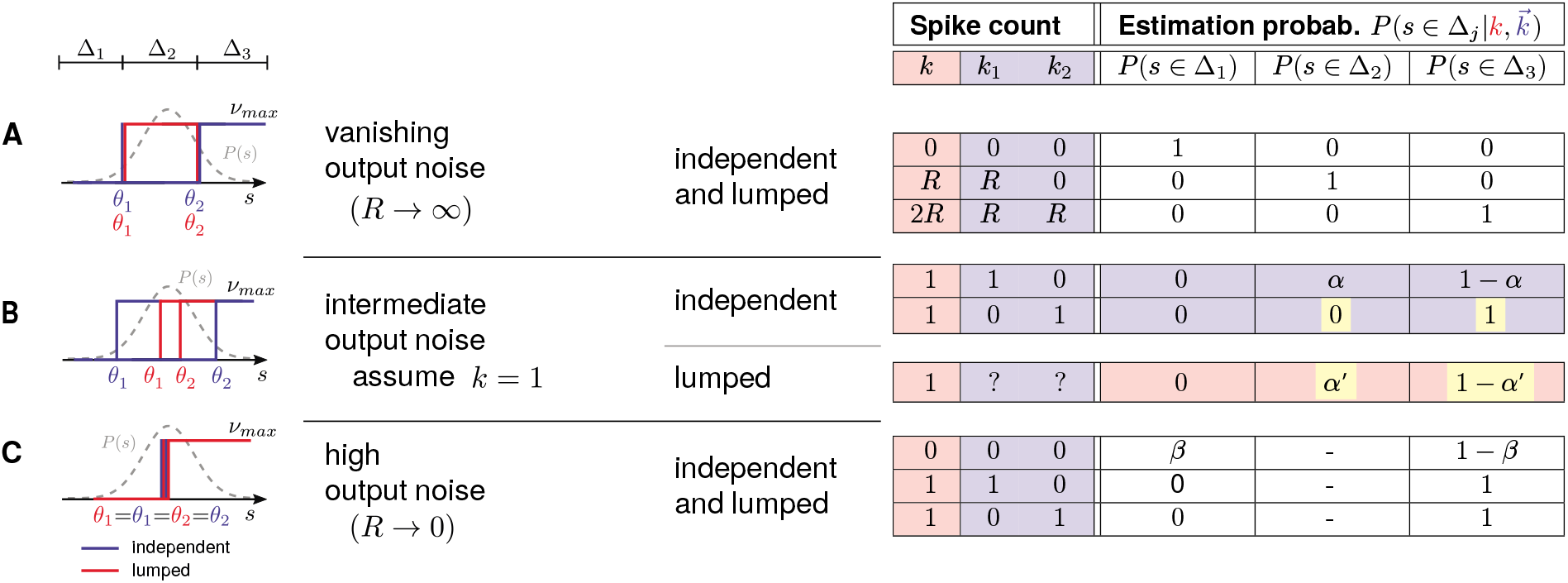
Schematic illustrating the dominance in information of the independent-over the lumped-coding channel. Here, we treat the case of *N* = 2 cells, vanishing input noise (*σ* = 0) and **A**. vanishing output noise (*R* → ∞), **B.** intermediate output noise when the total number of spikes *k* = *k*_1_ + *k*_2_ = 1, and **C.** high output noise (*R* → 0). Left: The relative positions of optimal thresholds of both the independent-(blue) and lumped-coding (red) channels. Right: The stimulus “estimation probabilities” 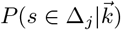 for the two different channels. Yellow shading shows where the noise entropy is higher in the lumped-coding channel. *α, α′*, and *β* denote non-zero probability values (see text).

For intermediate output noise, the independent- and the lumped-coding channels have distinct estimation probabilities. Although in principle the number of emitted spikes can be anything, let us consider the example where the total number of spikes is 1 (*k*_1_ + *k*_2_ = 1, Fig. 2B). We demonstrate that the lumped-coding channel loses information because knowledge about the identity of which individual cell spiked is lost. For example, if the cell with higher threshold *θ*_2_ fires at least one spike, this implies with certainty that the stimulus is greater than *θ*_2_. The lumped-coding channel fails to encode this information since in principle the spike could have been emitted by the cell with lower threshold *θ*_1_. Thus, the estimation probabilities *β*′ and 1 − *α*′ for the stimulus being below or above *θ*_2_, respectively, are nonzero. For the independent-coding channel, however, the corresponding estimation probabilities *α* and 1 − *α* are nonzero if the cell with the lower threshold θ1 fires a spike. Therefore, for the independent-coding channel there are more cases in which the uncertainty is resolved, leading to higher mutual information. As an example, for output noise of *R* = 2.5, the mutual information for the independent- and lumped-coding channel is 1.30 and 1.01 bits, respectively.

For very high output noise, *R* → 0, the expected spike count of either of the cells is very small, even when the stimulus is larger than the respective threshold with the resulting firing rate *ν*_max_ (Fig. 2C). This means that most of the time the observed spike count of each cell is 0, rarely 1, and never 2 – the probability of observing more than one spike is infinitesimal. In this high noise regime, the optimal solution for both the independent- and lumped-coding channels is to make both thresholds identical, i.e. *θ*_1_ = *θ*_2_. Therefore, the intermediate regime Δ_2_ does not exist and the two possibilities of having a spike from either cell are equivalent. Thus, if the observed spike count is 1, then there is no possibility of error for either channel. Similarly, if the observed spike count is 0, the two different estimation probabilities are the same for both channels, namely *β* and 1 − *β* for the stimulus being above or below *θ*_1,2_, respectively. This results in identical mutual information between stimulus and response for both channels.

Equipped with this intuition, we computed the mutual information between the lumped- and independent-coding channels for a population of three neurons where we varied both the input and output noise continuously (Fig. 3A,B). We again found that the independent-coding channel overall transmits more information than the lumped-coding channel. Additionally, we found that the increase of information with smaller output noise (higher *R*) saturates faster in the case of the independent-coding channel, as can be seen by the flattening of the contour lines as *R* increases (compare Fig. 3A and B).

**Figure 3.**
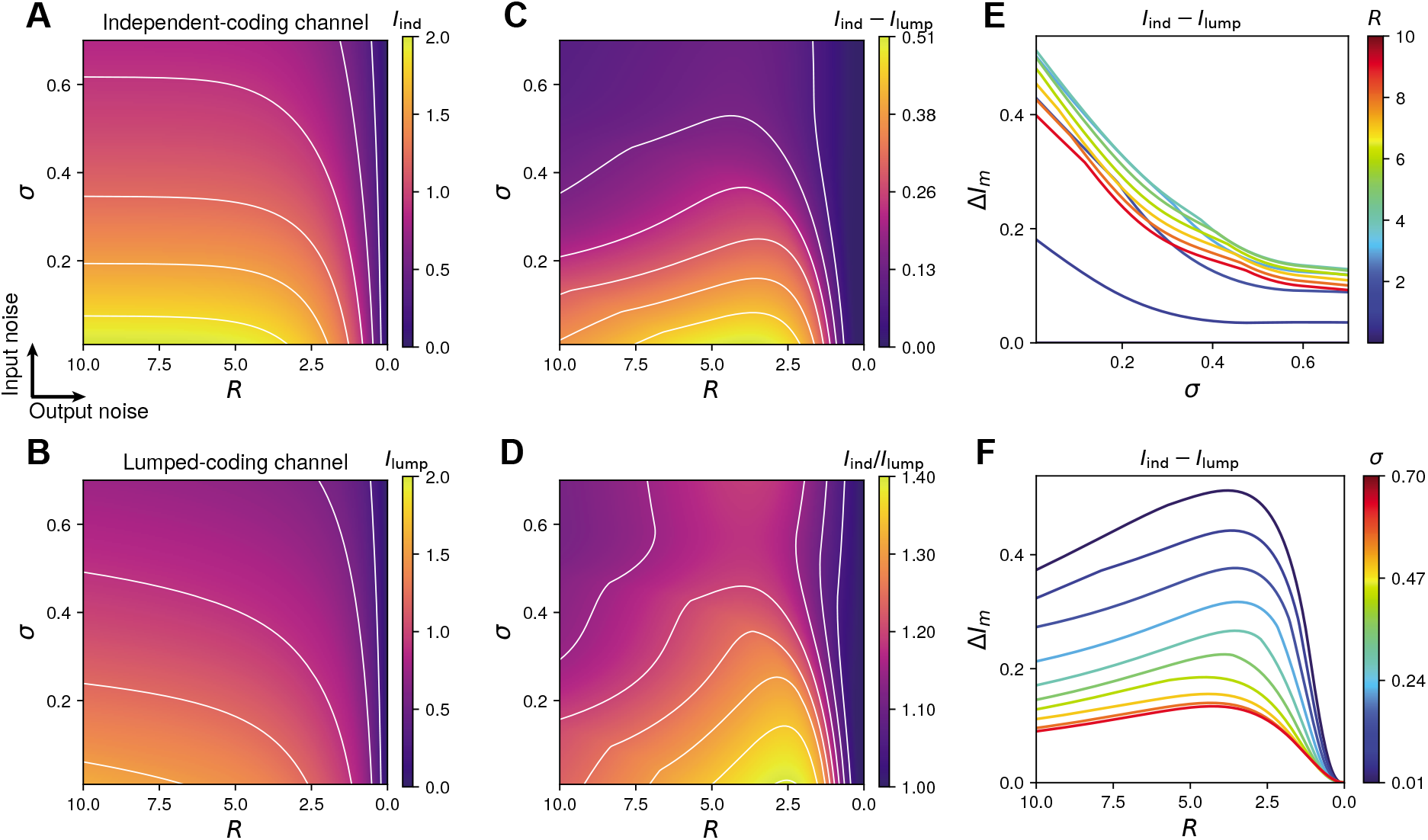
Mutual information for the lumped- and independent-coding channels for a population of three neurons. **A.** Information of the independent-coding channel for different combinations of output noise *R* and input noise *σ*. Contours indicate constant information. **B.** Information of the lumped-coding channel. **C.** Absolute information difference between the two coding channels. **D.** Information ratio between the two coding channels. Both C and D show a region of intermediate output noise where the independent-coding channel substantially outperforms the lumped-coding channel. **E.** Information difference depending on input noise **σ** for various levels of output noise *R*, corresponding to vertical slices from C. **F.** Information difference depending on output noise *R* for various levels of input noise *σ*, corresponding to horizontal slices from C.

We quantified the ratio and the absolute differences in information transmission between the two channels (Fig. 3C,D). For all finite input and output noise levels, the independent-coding channel outperforms the lumped-coding channel since the contribution of each neuron to the overall spike count provides additional information about the stimulus that is lost by summing all the spike counts through lumping. The information loss is the largest at intermediate levels of output noise and low levels of input noise; for instance, at *R* ≈ 2.5 and *σ* ≈ 0 the independent-coding channel transmits up to 40% more information than the lumped-coding channel (Fig. 3D).

To best visualize these differences, we fixed one source of noise and varied the other. For fixed output noise *R*, the information loss in the lumped-coding channel relative to the independent-coding channel monotonically decreases as a function of the input noise, *σ* (Fig. 3E). The difference in information transmitted by the independent- and lumped-coding channels as a function of the output noise *R* for fixed input noise *σ* demonstrates that the information loss due to lumping is a non-monotonic function of output noise *R* (Fig. 3F), with the largest loss occurring in the biologically realistic range of intermediate noise [36, 48]. This non-monotonicity can be explained by the fact that in the limit of very large or very small output noise the lumped-coding channel transmits as much information as the independent-coding channel (Fig. 2).

In summary, we found that in the presence of both input and output noise, the lumped-coding channel transmits less information than the independent-coding channel, and we can intuitively understand these trade-offs in a small population of two neurons.

### Optimal thresholds for the independent- and lumped-coding channels

We computed the optimal population thresholds at which the spiking output of the populations achieves maximal information about the stimulus. We first discuss the case with three neurons. For both the independent- and lumped-coding channel, the optimal number of distinct thresholds in the population depends on the source and level of noise (Fig. 4). When both sources of noise are negligible, the optimal number of thresholds is three, representing a fully diverse population where all thresholds are distinct. However, when both input and output noise are high, the optimal number of thresholds in the population is one, representing a full redundant population where all thresholds are identical. The most interesting cases arise at intermediate input and output noise levels, where we found two distinct optimal thresholds. To gain a better understanding of the transition between different threshold regimes as a function of noise, we fix one level of noise and examine the thresholds as a function of the other noise level.

**Figure 4.**
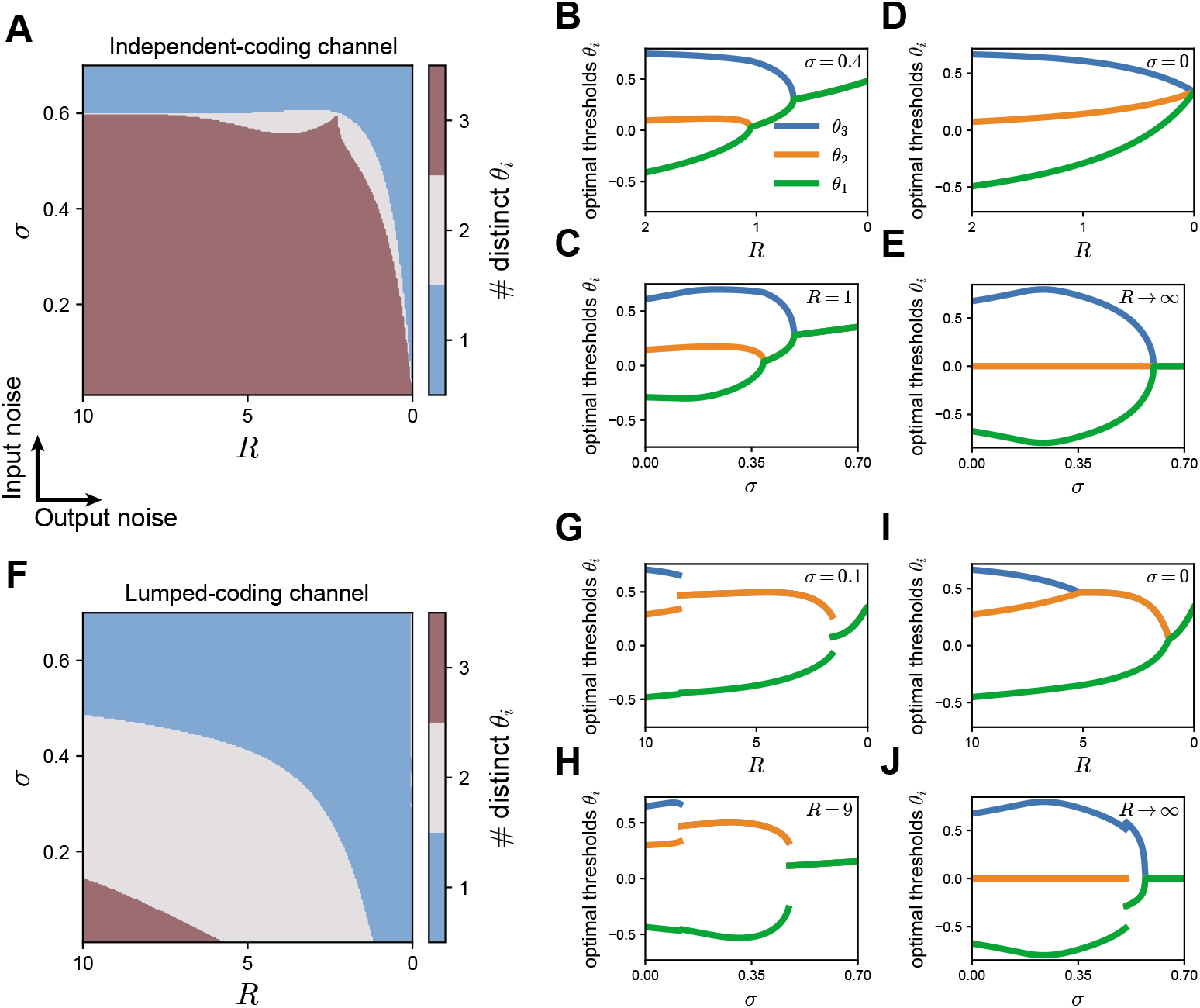
Optimal thresholds for the independent- and lumped-coding channels. Optimal thresholds for the independent-coding channel (A-E) is compared to the lumped-coding channel (F-J) for a population of three neurons. **A.** The optimal number of distinct thresholds depend on input noise *σ* and output noise *R*. **B.** The optimal thresholds as a function of output noise for a fixed value of input noise (*σ* = 0.4). **C.** The optimal thresholds as a function of input noise for a fixed value of output noise (*R* = 1). **D.** The optimal thresholds as a function of output noise in the limit of no input noise (*σ* = 0). **E.** The optimal thresholds as a function of input noise in the limit of vanishing output noise (*R* → ∞). **F-J.** As (A-E) but for the lumped-coding channel. Intermediate noise levels in (G,H) take smaller values of *R* and *σ* in the lumped-coding channel since lumping itself acts like a source of noise (G: *σ* = 0.1, H: *R* = 9).

We found that the number of distinct thresholds in the population generally decreases with increasing input or output noise through a set of bifurcations. We call the noise levels at which these bifurcations in the thresholds appear *critical* noise levels. We found that for the lumped-coding channel threshold bifurcations occur at lower noise levels compared to the independent-coding channel. This result makes intuitive sense because lumping multiple information pathways into a single coding channel reduces the possible values of the encoding variable and increases the noise entropy, and therefore acts like an additional noise source.

For the independent-coding channel (Fig. 4A), the thresholds become distinct from each other gradually, in the sense that the differences between the optimal thresholds change continuously, both as a function of output noise when the input noise level is fixed (Fig. 4B) and also as a function of input noise when the output noise level is fixed (Fig. 4C). In the case when one source of noise is zero, these bifurcations represent the transition from all optimal thresholds being distinct directly to the state where all optimal thresholds are identical, without an intermediate state where two thresholds are the same (Fig. 4D,E). For instance, in the absence of input noise (*σ* = 0), the population’s thresholds are all distinct from each other for all finite ranges of output noise except when *R* → 0 (Fig. 4D). In the absence of output noise (*R* → ∞), there is a critical value *σ*_crit_ > 0 at which the population transitions directly from all thresholds being distinct to all thresholds being equal (Fig. 4E). Note that for all these bifurcations the threshold differences change continuously, i.e. there are no jumps of optimal threshold values with varying noise.

Surprisingly, we found a small range of input noise, 0.54 < *σ* < 0.6, for which we observed a nonmonotonic change in the number of optimal thresholds when varying the output noise *R* (Fig. S1). A similar result has been observed when optimizing Fisher information – a different, local measure for information – in a population of bell-shaped tuning curves in a model of optimal coding of interaural time differences in the auditory brain stem [28].

In comparison, for the lumped-coding channel, the bifurcations occur as the threshold differences at critical noise values change abruptly, or discontinuously, when one noise source varies and the other remains fixed (Fig. 4G,H). Here, the system has an intermediate number of thresholds for a large range of noise values, and the transition from one to three distinct thresholds is not simultaneous as either noise vanishes. Rather, the discontinuous threshold jumps at each bifurcation become continuous (Fig. 4I), as normally seen for the independent-coding channel, or partly continuous (Fig. 4J). These two scenarios agree with two previous studies, where a lumped-coding channel was studied with only output noise (Fig. 4I) [30], or with only input noise (Fig. 4J) [29]. Our results are also consistent with previous studies for small populations of two neurons and only one source of noise [5,14,27], large populations with only output noise [39] and two-neuron populations with multiple noise sources [25]. We also show that similar patterns of how the number of distinct thresholds evolve as a function of two different noise sources for the independent-lumped-coding channels also hold for larger neural populations (Fig. S2).

Taken together, our theory derives different configurations of thresholds in populations of more than two noisy neurons that depend on how the sensory stimulus is combined to produce spiking output and the location of the noise that corrupts the signal.

### Optimal threshold differences represent order parameters in phase transitions

The characteristic bifurcations of the optimal thresholds at critical noise levels suggest the occurrence of phase transitions encountered in a variety of physical systems. In physics, a phase transition is characterized by the fact that an order parameter changes abruptly with external parameters [49]. For example, a phase transition occurs when the order parameter – which could among others be the density at the liquid-vapor critical point, or magnetization of a ferromagnetic material – changes abruptly with an external parameter, such as pressure or temperature. Similarly, in chemistry, the order parameter that changes abruptly with temperature is the solubility of liquid mixtures.

Guided by this characterization, we sought to relate the qualitative differences in optimal thresholds of the independent-vs. lumped-coding channel with two noise sources to phase transition phenomena (Fig. 4). We illustrate the results for a population with three neurons, and thus have two order parameters which are the two threshold differences, *θ*_1_ – *θ*_2_ and *θ*_3_ – *θ*_2_. To determine whether a phase transition occurs, we computed the first and second derivatives of the mutual information with respect to a given noise parameter (Fig. 5). Using the Ehrenfest classification of phase transitions [50], a discontinuity in the first (second) derivative with respect to the noise implies a first-(second-) order phase transition.

**Figure 5.**
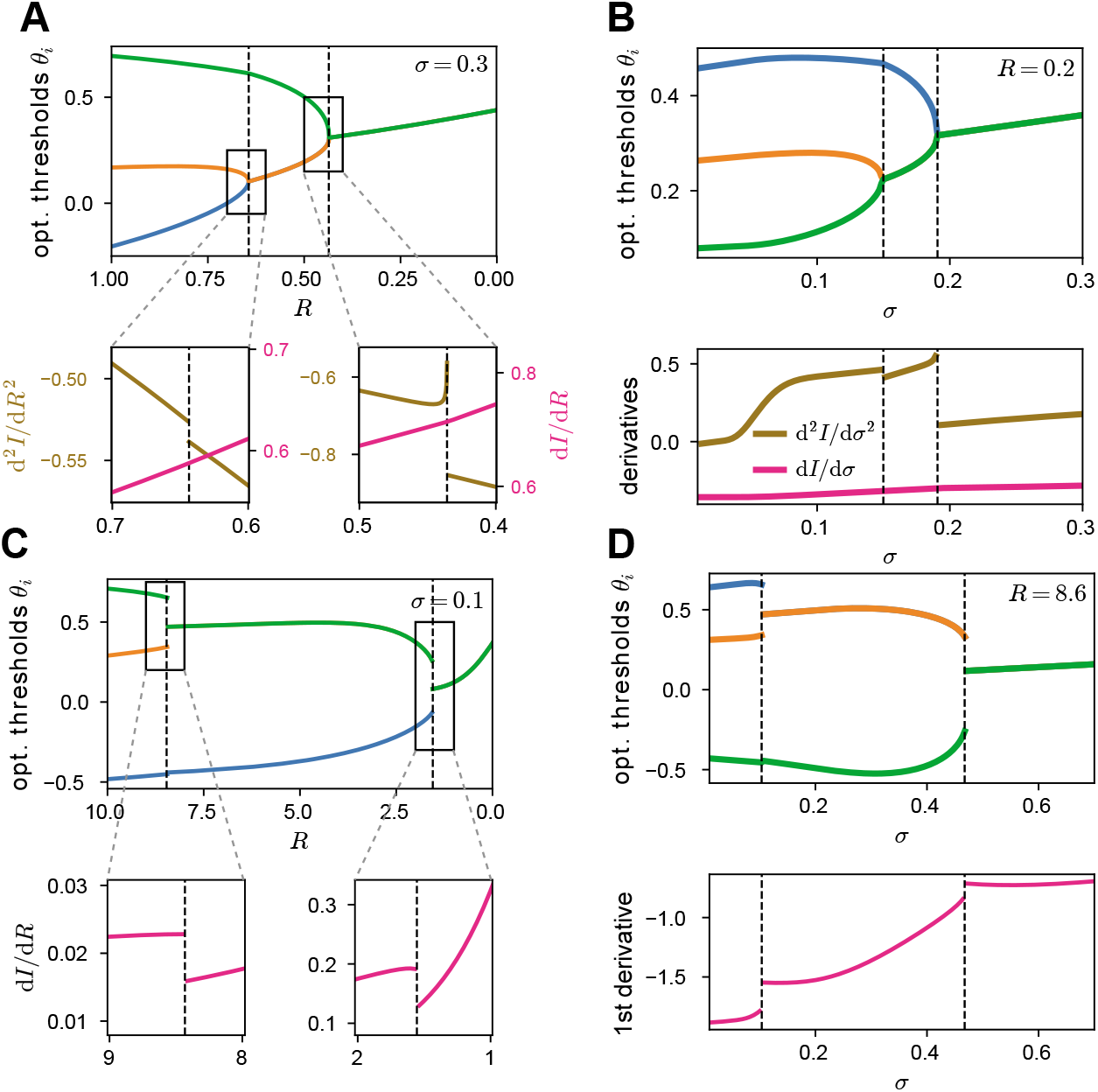
Threshold differences as phase transitions with respect to both noise sources. **A**. Optimal thresholds for the independent-coding channel depending on output noise *R*. Insets: The first derivative of the mutual information as a function of noise is continuous, while the second derivative is discontinuous at the critical noise values where the thresholds separate, implying a second-order phase transition. **B**. As in A, but with respect to input noise *σ*. **C**. Optimal thresholds as in A but for the lumped-coding channel. The first derivative is discontinuous at the critical noise values where the thresholds separate, implying a first-order phase transition. **D**. As in C but with respect to input noise *σ*.

We found that the orders of the phase transitions always correspond to the discontinuity of the threshold differences – being the order parameters – when noise varied. For continuous threshold bifurcations, there was a discontinuity in the second derivative with respect to output noise, thus corresponding to a second-order phase transition (Fig. 5A). All phase transitions for the independent-coding channel were continuous and thus of second-order also with respect to input noise (Fig. 5B). This result is in agreement with a previous study which also found a second-order phase transition in a population of two neurons in the presence of only input noise [5]; we extended this result to populations of more than two neurons and with additional output noise. We next investigated phase transitions in the lumped-coding channel.

For discontinuous threshold bifurcations we observed a discontinuity in the first derivative with respect to output noise and thus a phase transition of first-order (Fig. 5C). This is almost always the case for the lumped-coding channel, also with respect to input noise (Fig. 5D). An exception to this is when one noise source vanishes, e.g. input noise (Fig. 4I) or output noise (Fig. 4J), for which the phase transitions are of second-order (Fig. S3).

Together this shows that the continuity of the order parameter – here the threshold differences in the population of neurons – can be related to the order of the observed phase transition: discontinuous threshold differences correspond to first-order phase transitions, while continuous threshold differences correspond to second-order phase transitions.

### Characteristic shape of the information landscape at critical noise levels

To gain a better understanding of the information landscape, especially at the critical noise values at which threshold bifurcations appear, we examined the Hessian matrix of the mutual information, *I_m_*, with respect to the thresholds, *∂*^2^*I_m_*/(*∂θ_i_∂θ_j_*). The Hessian can be understood as an extension of the second derivative to higher-dimensional functions. The eigenvalues of the Hessian quantify the curvature of the information landscape in the direction of the respective eigenvectors, which themselves stand for the directions of principal curvatures in the space defined by the thresholds. To gain intuition about the differences of the information landscape between the independent- and lumped-coding channels, we considered a population of two cells for which the landscape can be easily portrayed in two dimensions. However, the theory extends naturally to populations with more neurons and information landscapes in higher dimensions.

We first considered the independent-coding channel for a fixed level of input noise, while varying the output noise. At the critical noise level, *R*_crit_, where the thresholds bifurcate, one eigenvalue of the Hessian decreases to zero (Fig. 6A). The information landscape undergoes a transformation around the critical noise levels, from one with two distinct maxima separated by a local minimum at low noise, *R* > *R*_crit_ (Fig. 6B, top), where the population thresholds are distinct, to one where there is a unique maximum at high noise, *R* < *R*_crit_, where the population thresholds are identical (Fig. 6B, bottom). For *R* > *R*_crit_, there are two inflection points (Fig. 6C, top), resulting in two different curvatures along the line that connects the two maxima. At the critical noise, *R* = *R*_crit_, the two maxima converge at the bifurcation point and the two inflection points fuse together such that the curvature becomes zero (Fig. 6C, middle). At this point of convergence, the information landscape locally resembles a ridge, which extends along one principal direction of curvature (Fig. 6B, middle). The ridge is perpendicular to the other principal direction, which stands for the direction of largest curvature. Finally, for *R* < *R*_crit_, the information landscape has a single maximum with negative curvature (Fig. 6B and C, bottom).

**Figure 6.**
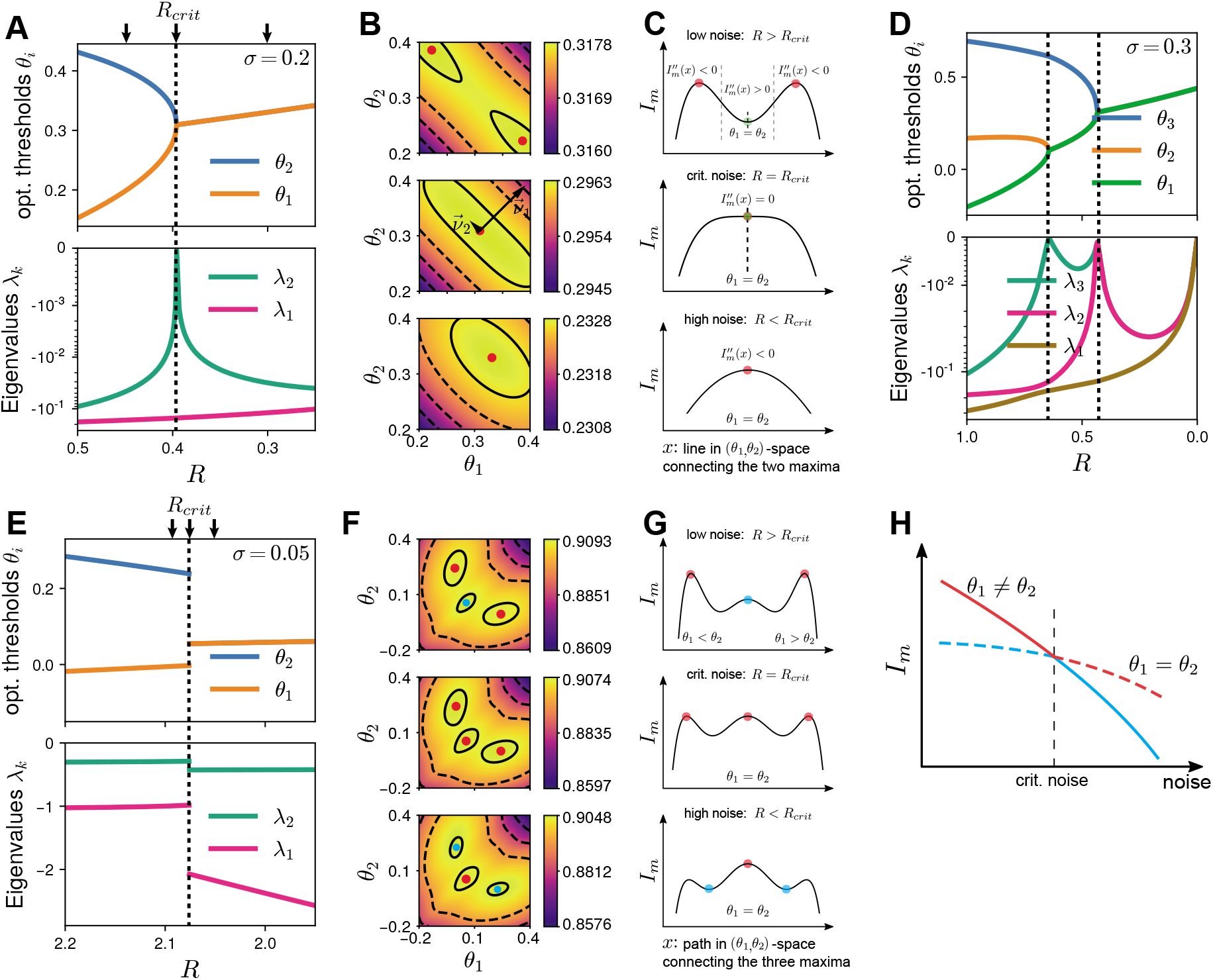
Information landscape for the independent- and lumped-coding channels undergoes different phase transitions at critical noise levels. **A**. Top: Optimal thresholds of the independent-coding channel for a population of two neurons as a function of output noise *R*. Bottom: Corresponding eigenvalues of the Hessian of the information landscape with respect to thresholds. At the critical noise value *R*_crit_ ≈ 0.396 at which the threshold bifurcation occurs (vertical dashed line) the smaller eigenvalue approaches zero. **B**. Information landscape *I_m_*(*θ*_1_, *θ*_2_) for the three output noise levels *R* indicated by arrows in A. Top: For *R* > *R*_crit_, there are two equal maxima. Middle: At *R* = *R*_crit_, the eigenvectors of the Hessian are shown and scaled by the corresponding eigenvalue (the eigenvector with the smaller eigenvalue, 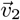, was artificially lengthened to show its direction). At the critical noise value the information landscape locally takes the form of a ridge. Bottom: For *R* ≤ *R*_crit_, there is one maximum, meaning that the optimal thresholds are equal (bottom). **C**. The mutual information as a function of the line *x* in (*θ*_1_, *θ*_2_) space connecting the two maxima in B. Top: For *R* > *R*_crit_ (low noise), there are two inflection points (dashed vertical lines) with zero curvature along the line *x*. The point with equal thresholds corresponding to a local minimum. Middle: At *R* = *R*_crit_, the two maxima, the minimum, and the two inflection points merge in one point, thus the curvature is zero. Bottom: For *R* ≤ *R*_crit_, there is a single maximum with negative curvature. **D**. As in A but for a population with three neurons. **E**. As in A but for the lumped-coding channel. Both the optimal thresholds and the eigenvalues show a discontinuity at critical noise level. **F**. Information landscape as in B for lumped-coding channel and noise values indicated by arrows in E. Local maxima are shown in cyan, global ones in red. **G**. Similar to C for lumped-coding channel. Here the abscissa denotes the (non-straight) path connecting the three maxima in F. **H**. Illustration of discontinuous threshold bifurcations, where the global maximum at *θ*_1_ ≠ *θ*_2_ at low noise (red, solid) becomes a local maximum for high noise (cyan, solid), while *θ*_1_ = *θ*_2_ (dashed) becomes global. As their respective derivatives are different, there is a discontinuity in the first derivative when only considering the global maximum (red lines), corresponding to a first-order phase transition.

We then examined the eigenvalues of the Hessian matrix for a larger population of size *N*. We found that at each critical noise level where the thresholds bifurcate, at least one eigenvalue of the Hessian matrix approaches zero. The number of zero eigenvalues – denoting the number of dimensions along which the information does not change locally – is equal to the number of thresholds participating in a bifurcation minus one. For *N* = 3, for example, there are two critical noise values at which the thresholds bifurcate (Fig. 6D, top). At one of these critical values, three thresholds are involved and thus the number of eigenvalues approaching zero is two, while at the other critical value only two thresholds are involved, and thus the number of eigenvalues approaching zero is one (Fig. 6D, bottom). Locally, threshold combinations along the ridge of the information landscape achieve almost the same information. This ridge is a manifold of dimension *M* − 1, where *M* is the number of thresholds involved in the bifurcation. The manifold is locally given by

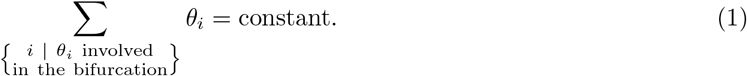

As a result, the ridge is oriented at exactly 45° with respect to all of the *θ*-directions participating in the bifurcation. For example, for *M* = 2 this manifold is a line, while for *M* = 3 it is a plane. Following the same argument as for the population with *N* = 2 neurons (Fig. 6C), it can be shown that the curvature of the information landscape has to be zero in *M* – 1 principal directions, thus *M* – 1 eigenvalues of the Hessian have to be zero when *M* thresholds participate in a bifurcation of continuous manner.

For the lumped-coding channel, the eigenvalues of the Hessian do not approach zero at the critical noise levels where the thresholds split (Fig. 6E). This is in agreement with the fact that threshold bifurcations are in general discontinuous for the lumped-coding channel (see also Fig. 4G,H). An exception to this is the limiting case when one noise level is zero, where the lumped-coding channel shows continuous bifurcations and thus second-order phase transitions (Fig. S3). At low output noise, *R* > *R*_crit_ (Fig. 6F,G, top), the information landscape has two distinct maxima corresponding to the optimal thresholds, *θ*_1_ and *θ*_2_. However, the information landscape also has a local maximum at *θ*_1_ = *θ*_2_. As noise increases, this local maximum decreases more slowly compared to the two global maxima, until at the critical noise level *R*_crit_ the three maxima become equal (Fig. 6F,G, middle). As noise increases further, *R* < *R*_crit_, the maximum at *θ*_1_ = *θ*_2_ becomes the single global maximum (Fig. 6F,G, bottom). Therefore, the phase transition happens at the noise level where the local maximum becomes the global one. This is a first-order phase transition since at this critical noise level the decrease of maximum information with noise changes abruptly, resulting in a discontinuity in the first derivative (Fig. 6H).

Our results show, that for finite noise, the shape of the information landscape for the independent- and the lumped-coding channels can be uniquely related to the nature of the threshold bifurcations (continuous for the independent-coding and discontinuous for the lumped-coding channel), and thus to the order of the phase transition. The information landscape takes a qualitatively different shape at the threshold bifurcations in each case, demonstrating the emergence of a new threshold through splitting either through a gradual “breaking” of the information ridge (Fig. 6B,C), or through a discrete switching from a local information maximum to the global maximum (Fig. 6F-H).

### Thresholds in auditory nerve fiber population match predictions from optimal coding

Next, we sought to compare our theoretical predictions of optimal thresholds to experimentally recorded thresholds of sensory populations to determine whether they are consistent with optimal coding. Specifically, we considered recordings of auditory nerve fibers (ANFs) coding for sound frequency and sound intensity. At the first synapse level of the auditory pathway, each inner hair cell of the cochlea transmits information about sound intensity to approximately ten to thirty different ANFs [7]. ANFs differ in several aspects of their responses, including spontaneous rates and thresholds, with each ANF receiving input exclusively from only a single inner hair cell [8]. We investigated the properties of experimentally recorded ANF tuning curves in the mouse [8]. As a population, ANF tuning curves resemble a sigmoid which increases with sound intensity; the sigmoid can be described by a threshold, a dynamic coding range also referred to as the gain, a spontaneous firing rate and a maximal firing rate (see Methods, Fig. 7A). Interestingly, ANF response curves with higher spontaneous firing rate have been shown to have narrower dynamic ranges and higher thresholds [8] (Fig. 7B). Given the lack of convergence on stimulus channels, we investigated whether our theoretical framework of independent-coding (rather than the lumped-coding) channel with two sources of noise before and after a nonlinearity can be applied to explain this relationship between spontaneous firing rate, dynamic range and firing threshold, testing the hypothesis that ANFs have optimized their response properties to encode maximal information about the stimulus under biological constraints.

**Figure 7.**
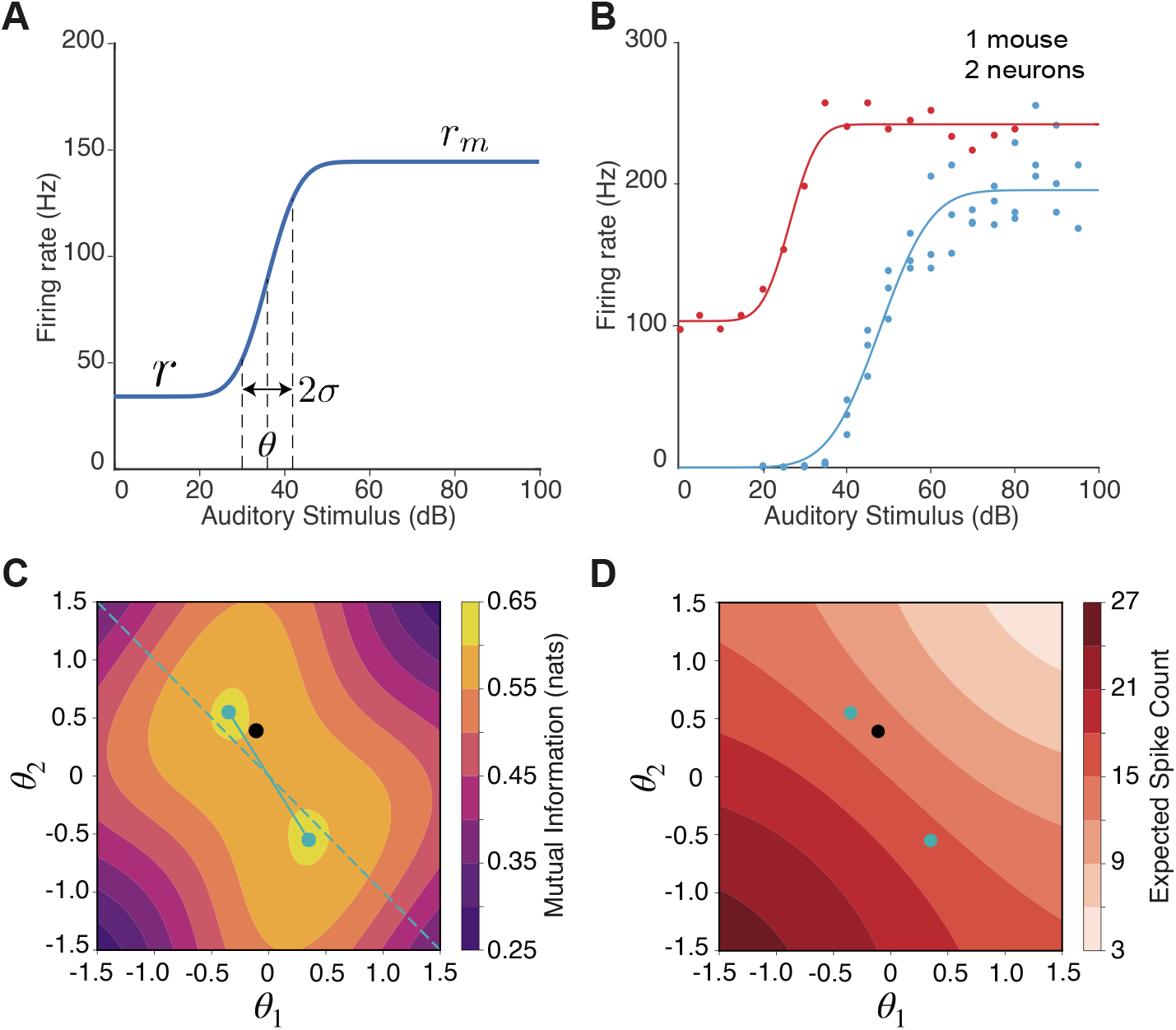
The tuning of auditory nerve fibers (ANFs) match predictions from optimal coding. **A.** The sigmoidal function that we use to fit ANFs tuning curves. Spontaneous firing rate (*r*), maximal firing rate (*r_m_*), firing threshold (*θ*), and the dynamic range (*σ*) are labelled on the curve. **B.** An example showing original data from ref. [8] and fitted tuning curves. These two tuning curves come from the same mouse. **C.** Optimal configuration (cyan dots) in the contour plot of mutual information. Black dot denotes the fitted thresholds from the data. **D.** Optimal configuration (cyan dots) in the contour plot of average firing rate. Black dot denotes the fitted thresholds from the data.

To apply our population coding framework to this type of data would require measurements of the entire population of ANFs. In the absence of such data, we decided to apply the framework to a population of two representative neurons where each neuron can be described by a sigmoidal response function, one with a high and the other with a lower spontaneous firing rate. This is computationally tractable and consistent with previous literature [8,51]. To obtain the two representative neurons, we proceeded as follows: first, we pooled the measured tuning curves from the same ANF, and fitted each with our sigmoidal function (see Methods, Fig. 7A). This resulted in 148 tuning curves from 24 animals. Indeed, we confirmed that the dynamic range is negatively correlated with the normalized spontaneous firing rate, as well as positively correlated with the thresholds (Fig. S4). Then, we divided all the tuning curves into two types based on their normalized spontaneous firing rate and dynamic range (see Methods) [8,51]. In particular, the ‘Type 1’ neuron had a higher spontaneous rate, smaller dynamic range and a lower firing threshold, while the ‘Type 2’ neuron had a lower spontaneous rate, broader dynamic range and a higher firing threshold (Fig. 7B, Fig. S4).

For this two-neuron population where one neuron had a nonzero and the other zero spontaneous rate, we optimized the neuronal thresholds while maximizing the mutual information between the stimulus and the population response. We used the level of input noise to modulate the neurons’ dynamic range (see Methods). Using dynamic ranges for the two neurons chosen to match the two neuron types found in the data, we evaluated the mutual information between stimulus and response for a range of firing thresholds (*θ*_1_ and *θ*_2_). We found that the mutual information landscape is centrosymmetric, with a maximal value of 0.603 nats achieved for two pairs of thresholds: (*θ*_1_, *θ*_2_) = {(−0.35, 0.55), (0.35, −0.55)} (Fig. 7C, cyan symbols). However, the second pair yields ~ 20% higher mean firing rate (Fig. 7D, cyan symbols). Therefore, the information per spike was higher for the first pair (0.0436 vs. 0.0378 nats/spike). When we overlaid the fitted thresholds from the data, we found that they lie remarkably close to the optimal thresholds obtained from the theoretical analysis (Fig. 7C,D, black symbol), suggesting that the ANF population might be configured to maximize information per spike about a distribution of frequencies.

We further explored how sensitive the optimization is to the chosen parameters of the sigmoids of the two types of neurons, more specifically, the dynamic range and the spontaneous rate. When both neurons are binary (no input noise) so that the sigmoids are infinitely steep and have a narrow dynamic range, and the spontaneous rate in both neurons is zero, the contour of the mutual information is both axisymmetric and centrosymmetric but the two pairs of optimal thresholds yield identical mean firing rate (Fig. S5A). Even when the neurons acquire a finite gain (by increasing input noise) so that the dynamic range is broadened, as long as the gain is the same, the information per spike for the two pairs of optimal thresholds remains identical (Fig. S5B). Either changing the spontaneous rate or the gain of one of the neurons can break the symmetry in the information landscape, such that it loses its axisymmetry but preserves its centrosymmetry. The symmetry breaking effect is much stronger when the gains of the two neuron types are different (Fig. S5C vs. D), and combining both different gain and non-zero spontaneous rate as in the data preserves the strong symmetry breaking (Fig. S5E-G). Intuitively, the neuron with the larger input noise and hence, lower gain, has a threshold with a larger absolute value so that it is farther away from the mean of the stimulus distribution and is thus less affected by the input noise (Fig. S5H), just like what we found in the data. Therefore, our simple theoretical model with binary neurons and two sources of noise can explain the relationship between spontaneous rate and thresholds in two types of ANFs in the mouse by maximizing information per spike.

## Discussion

We maximized mutual information between stimulus and responses of a population of neurons which encode a one-dimensional stimulus with a binary nonlinearity corrupted by two different noise sources, specifically, additive input noise before the nonlinearity and Poisson output noise after the nonlinearity. We compared two frameworks for stimulus convergence commonly used in previous studies, specifically, encoding the stimulus with independent transmission channels [5, 14, 25, 39] or lumping the channels into one effective channel [29,30]. In each scenario, we calculated the optimal thresholds of the population (Fig. 4).

### Lumping of information channels as a coding strategy with low cost

Unsurprisingly, increasing either input or output noise in the population, decreases the total amount of transmitted information; but the independent-coding channel always encodes more information than the lumped-coding channel, especially for biologically realistic, intermediate output noise values (Fig. 3). This occurs because lumping multiple information pathways into a single coding channel reduces the possible values of the encoding variable and increases the noise entropy, thus introducing additional noise. Therefore, threshold bifurcations in the lumped-coding channel occur at significantly lower critical noise levels compared to the independent-coding channel (Fig. 4).

Why would a biological system lump information transmission channels? A biological upside of combining information from multiple streams into one effective channel could be the reduction of neurons needed for information transmission, thus saving space and energy. For example, the optic nerve has a strong incentive to reduce its total diameter since it crosses through the retina and thus causes a blind spot. On the other hand, for a given constraint on space and energy, it is favorable to have many thin, low-rate axons over fewer thick, high-rate axons [52,53], thus arguing against convergence. However, at least for the retina, an intermediate degree of convergence is probably the optimal solution. One would expect that this degree of convergence depends on the location at the retina. At the fovea of a primate retina, there is minimal convergence from photoreceptors to retinal ganglion cells compared to the periphery [54]. This implies that a higher visual acuity is achieved by increasing information transmission at the cost of energy and space. In contrast, there does not seem to be any convergence in the early auditory pathway: At the first stage of the neural signaling process, one inner hair cell diverges to 10 to 35 auditory nerve fibers [7]. This lack of convergence might be due to the fact that, contrary to the retina, there is no pressure of having a thin ganglion. A recent theoretical study suggests that convergence can compensate the information loss due to a nonlinear tuning curve with a small number of output states [55].

We only treated the extreme cases of full convergence – where all neurons are lumped into a single channel – and no convergence. In principle, different combinations of partial convergence, e.g. lumping three outputs into two channels, are also possible. Partial lumping is a common strategy in sensory systems with different levels of convergence [56].

### Optimal number of distinct thresholds as a function of noise

The number of distinct optimal thresholds decreases with increasing noise of either kind at critical noise levels by successive bifurcations of the optimal thresholds (Fig. 4). We mapped these characteristic bifurcations of the optimal thresholds at critical noise levels to phase transitions of different orders with order parameters being the threshold differences. At finite noise levels, the lumped-coding channel undergoes discontinuous threshold bifurcations which correspond to a first-order phase transition with respect to noise where the threshold differences are the order parameters. In contrast, for the independent-coding channel, the threshold differences change continuously and the phase transitions are of second-order.

Our results suggest that input and output noise influence the mutual information in the same way that temperature affects free energy in physical systems [49]; in that sense, both noise sources act as external parameters with respect to which the phase transition occurs. In contrast to most physical systems, we have more than one order parameter, specifically the number of subsequent threshold differences, i.e. the number of neurons minus one. Our scenario with three neurons shows similarities with a system with three mixed liquids where the miscibility depends on the liquids’ relative concentration differences [57]. As the temperature varies, the system undergoes phase transitions where the miscibility changes, from having one phase in which all three liquids are mixable (similar to our scenario with three identical thresholds), to two phases where in one phase two liquids are mixable but which is separated from a second phase containing the third liquid (corresponding to two distinct thresholds in our neural population), to three phases where none of the liquids are mixable with each other (corresponding to the case of all distinct thresholds). As in classical physical systems, the orders of our phase transitions are consistently linked to the continuity of the threshold differences: a continuous (discontinuous) order parameter corresponds to a second (first) order phase transition.

Interestingly, for a range of noise parameters, we found a non-monotonic change in the number of distinct optimal thresholds with noise levels (Fig. S1). A similar non-monotonicity has also been reported under maximization of the Fisher information for neurons encoding sound direction [28]. This happens because of how the different neurons tile their thresholds to optimally encode the one-dimensional stimulus in the presence of multiple noise sources which interact non-trivially. The biological implications of such a non-monotonic change in the number of optimal thresholds as a function of noise are unclear. A related phenomenon in physics is that of *retrograde phenomena* [58]. For example, in a mixture of liquids, a phase transition from liquid to gas, followed by another transition from gas to liquid, and then liquid to gas again can be observed while increasing temperature [58].

### Information loss at non-optimal thresholds

An important, but often neglected, question for optimal coding theories is how much worse are suboptimal solutions in comparison to optimal ones in terms of information transmission. In the independent-coding channel, near critical noise levels, the information landscape becomes flat in the directions of principal curvature. This suggests that multiple threshold combinations yield nearly identical information, a property of the neural population that is closely related to the concept of ‘sloppiness’ whereby a system’s output is insensitive to changes in many parameter combinations but very sensitive to a few [59–61]. Hence, even population codes that utilize suboptimal thresholds often achieve information very close to the maximal, and it is unclear whether such small information differences could be measured experimentally. This also raises the question whether a few percent more information about a stimulus realized by optimal codes could be sufficiently beneficial for the performance of a sensory system to become a driving force during evolution. It has been shown that mutations which have very small effects on evolutionary fitness are fixated in a population with a probability almost irrespective of the mutation being advantageous or deleterious [62, 63]. On the other hand, in certain sensory systems like the retina, entire populations of retinal ganglion cells perform multiple functions [64,65] or fulfill different computations under different light conditions [66]. For such systems, there must be a fundamental trade-off in performance, since such a system cannot be optimal at all functions [67, 68]. The sloppiness of nearly-equivalent optimal thresholds that we observe near critical noise levels should resolve when considering that neurons have multiple constraints and often perform more than just one function or encode different stimulus features.

### Assumptions in our model and comparison to other theoretical frameworks

There are several modeling assumptions in our theoretical framework that make mathematical treatment possible. First, we considered the encoding of a static stimulus, even though natural stimuli have correlations in space and time. Previous studies have exploited their correlation structure to explain various aspects of sensory coding, for example, the size and shape of receptive fields of retinal ganglion cells [12,13,21,33,35,38]. Since correlations in the stimulus are thought to reduce effective noise values [38], by considering stimuli independent in time, we likely underestimated effective noise levels.

Moreover, our coding framework assumed a one-dimensional stimulus; thus, it is appropriate for explaining the number of the population’s distinct thresholds which encode a *single* stimulus feature – this could be the contrast at a single spatial position on the retina (as found to be coded by two different types of OFF retinal ganglion cells that encode the same linearly filtered stimulus [5]), or sound intensity at a single frequency (as found to be coded by ANFs, which get input from the same inner hair cell [8, 69]). Throughout this study we investigated the encoding of a one-dimensional stimulus drawn from a Gaussian distribution; however, natural stimulus distributions have a higher level of sparseness than the Gaussian distribution [70,71]. Therefore, we also explored information maximization using the generalized normal distribution allowing us to continuously vary the kurtosis – how heavy the tails are – of both the stimulus and the input noise distributions. Our results remain qualitatively the same as for the Gaussian distributions (Fig. S6).

Second, we modeled each neuron in the population solely with a binary nonlinearity. This nonlinearity describes the tuning curve of the neuron as a function of a given stimulus feature. In general, a tuning curve with respect to a stimulus feature is measured by reverse correlating the stimulus variable with the output variable and fitting a linear-nonlinear model [72]. The linear part of the model denotes the stimulus feature to which the neuron responds and the nonlinear part represents the tuning curve. We did not incorporate the linear part in our model but rather assumed that the input to the nonlinearity is already linearly preprocessed because simultaneous optimization under different noise sources and stimulus convergence would be mathematically intractable. We chose binary nonlinearities as they are theoretically optimal under certain conditions of high (and biologically plausible) Poisson noise [25, 30, 73]. Importantly, however, under conditions of non-negligible input noise the optimal nonlinearity could be interpreted to acquire a finite slope thus making our analysis relevant also for continuous nonlinearities with sigmoidal shape. This is consistent with neuronal recordings; for example the steepness of the tuning curve of the H1 blowfly neuron increases with contrast, and for high contrast – which corresponds to low noise – the tuning curve is almost binary [16].

Third, we considered a constraint on the maximum expected spike count since the total encoded information cannot be infinite. Such a constraint is motivated by a biophysical limit of a neuron’s firing rate and the biological reality of a short reaction time. Instead, one could constrain the *mean* spike count [5,14,26], which would be interpreted as a metabolic constraint. Maximum and mean rate constraints lead to qualitatively similar conclusions regarding the optimal number thresholds, as shown in small populations of two neurons [5,14].

Many previous studies make very similar assumptions but consider certain limiting scenarios, for instance considering only one noise source [5, 29, 30, 39], studying a population with only two neurons [5, 25, 39], or introducing an additional source of additive output noise [25]. Table 1 summarizes these studies with regards to the different optimization measures, constraints, information convergence strategies, sources of noise and neural population size. While our results are in agreement with these previous studies in the specific limiting conditions, we extend the optimal coding framework by mapping the full space of noise and stimulus convergence thus linking and extending previous findings.

**Table 1.**
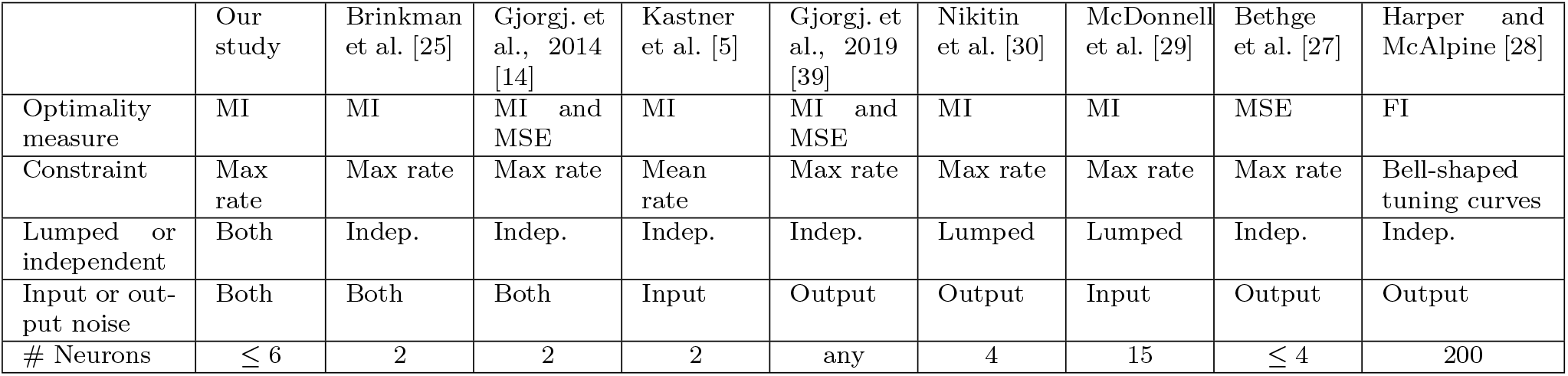
Comparison of different studies with regards to the different optimization measures, constraints, information convergence strategies, sources of noise and neural population size. MI stands for Mutual Information, FI for Fisher Information and MSE for Mean Square Error.

|  | Our study | Brinkman et al. [25] | Gjorgj. et al., 2014 [14] | Kastner et al. [5] | Gjorgj. et al., 2019 [39] | Nikitin et al. [30] | McDonnell et al. [29] | Bethge et al. [27] | Harper and McAlpine [28] |
| --- | --- | --- | --- | --- | --- | --- | --- | --- | --- |
| Optimality measure | MI | MI | MI and MSE | MI | MI and MSE | MI | MI | MSE | FI |
| Constraint | Max rate | Max rate | Max rate | Mean rate | Max rate | Max rate | Max rate | Max rate | Bell-shaped tuning curves |
| Lumped or independent | Both | Indep. | Indep. | Indep. | Indep. | Lumped | Lumped | Indep. | Indep. |
| Input or output noise | Both | Both | Both | Input | Output | Output | Input | Output | Output |
| # Neurons | $\leq 6$ | 2 | 2 | 2 | any | 4 | 15 | $\leq 4$ | 200 |

## Conclusions

In sum, we considered an optimal coding framework with contributions from two sources of noise and investigated information transmission under two different scenarios of stimulus convergence. Since we did not model a specific sensory system, but rather aimed to uncover general principles of optimal coding solutions under the two sets of independent scenarios above (noise and stimulus convergence), the sources of noise in our model do not directly correspond to circuit elements, making direct comparison to experimental data difficult. However, by applying our framework to coding by two types of ANFs in the mouse with a higher and lower spontaneous rate, we found that their thresholds are close to the optimal ones when maximizing information per spike. More importantly, we extended previous theoretical results that considered specific limiting scenarios, in the process providing a unifying framework for how different noise sources and the strategy of stimulus convergence influence information transmission and number of distinct thresholds in populations of nonlinear neurons.

## Methods

We assume that the stimulus *s* follows a Gaussian distribution with mean zero and variance 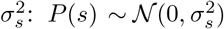. It is encoded by the spike counts {*k_i_*} of *N* binary neurons *i* = {1,..,*N*} in a given coding time window Δ*T*. Input noise *z*, 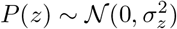, is added to the stimulus before the nonlinear processing, *σ*:= *σ_z_*/*σ_s_* denotes the effective amount of input noise. For both the stimulus and the noise distribution, we also considered other distributions with different kurtosis. However, we did not find significant differences to the Gaussian distributions (Fig. S6). We assume *N* binary nonlinearities *ν_i_*(*x*) = *ν*_max_Θ(*x* – *θ_i_*) with the two firing rate levels *ν_i_* = {0, *ν*_max_} and a respective threshold *θ_i_*. The input to each nonlinearity is the sum of stimulus and input noise: *x* = *s* + *z*. Poisson output noise is implemented by assuming that the spike count *k_i_* in the coding window Δ*T* follows the Poisson distribution, 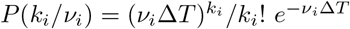.

We denote the expected spike count of a neuron firing with its maximum firing rate *ν*_max_ by *R*:= *ν*_max_Δ*T*. If *R* is small it means more output noise since even in the presence of maximum firing rate there are more occurrences of zero spikes and thus there is a higher ambiguity about the real firing rate. The above implementation of output noise can be understood as a constraint on the maximum firing rate level *ν*_max_ while having a fixed coding window length Δ*T*.

For given noise levels *σ* and *R* the goal is to find nonlinearities which optimally encode the stimulus *s* with a vector of spike counts 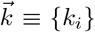 (independent-coding channel) or the lumped spike count 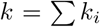 (lumped-coding channel). Since we assume binary nonlinearities and keep the two firing rates fixed, the only variables to optimize are the components of the threshold vector 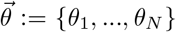. As a measure for optimality for the independent- and lumped-coding channels we choose the mutual information between stimulus *s* (input) and observed spike count 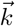 or *k*, respectively (output). The mutual information gives an upper bound on how much information can on average be obtained about the input by observing the output. It is given as the difference between output entropy 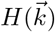 and noise entropy 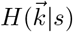 [74]:

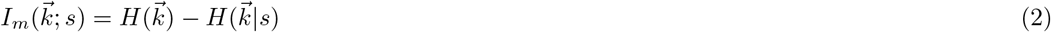

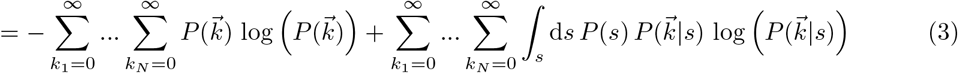

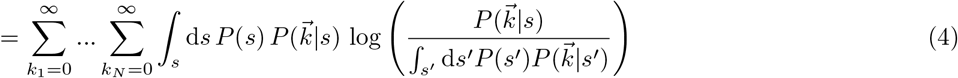

where the input-output kernel 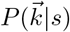 is the probability of obtaining a certain vector of output spikes for a given stimulus value. In the case of the lumped-coding channel the calculations are the same, except that the spike count is now one-dimensional, i.e. we have *I_m_*(*k*; *s*) as the mutual information and *P*(*k|s*) as the input-output kernel.

### Independent-coding channel

In the case of the independent-coding channel, 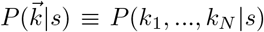 can be expressed by multiplying *P*(*k*_1_,…, *k_N_*|*ν*_1_,…, *ν_N_*) and *P*(*ν*_1_,…, *ν_N_*|*s*) and summing over all possible firing rate states {0, *ν*_max_}:

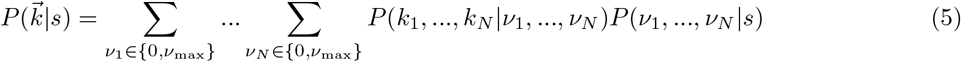

We assume no noise correlations and thus *ν_i_* conditional on *s* are independent of each other:

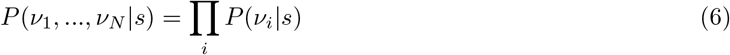

Furthermore, all *k_i_* are independent of each other conditional on a set of firing rates {*ν*_1_,…, *ν_N_*}, and every *k_i_* only depends on *ν*_*j*=*i*_:

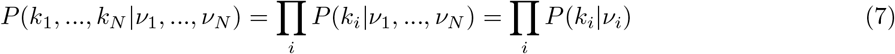

Taken together:

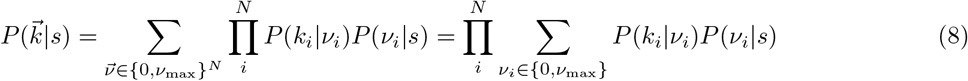

*P*(*k_i_*|*ν_i_*) follows a Poisson distribution and *P*(*ν_i_|s*) denotes the probability of having a firing rate of zero (or *ν*_max_) for a given stimulus *s*. Since the input noise fluctuations are on a much faster time scale than the length of the coding window (over which the stimulus is assumed to be constant), an averaging over *z* can be performed. Thus *P*(*ν_i_* = 0|*s*) (or *P*(*ν_i_* = *ν*_max_|*s*)) is given as the probability that stimulus plus noise is smaller (or larger, respectively) than threshold *θ_i_*, which is the area under the noise distribution for which *s* + *z* < *θ_i_* (or *s* + *z* ≥ *θ_i_*, respectively):

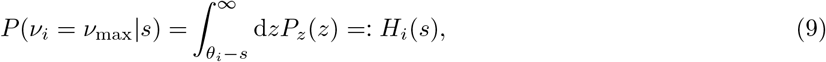

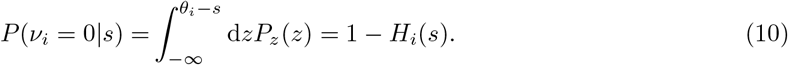

*H_i_*(*s*) can be viewed as the “effective” tuning curve that one would measure electrophysiologcally (see also Fig. 1, top right). It is the cumulative of the dichotomized noise distribution. If the noise distribution is normal distributed with variance *σ*^2^, the effective tuning curve is given by the complementary error function:

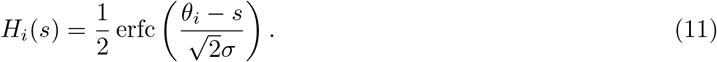

Then one can calculate the mutual information by performing the summation over all output variables *k*_1_,…,*k_N_*. The output noise is included since *P*(*k_i_|ν_i_*) is Poisson distributed. According to the Poisson distribution, *P*(*k_i_* = 0|*ν_i_* = *ν*_max_) = *e*^−*R*^. For each *k_i_*, all spike counts greater than zero can be lumped into one state due to the fact that if there is one or more spikes emitted, the firing rate can not be zero, i.e. *P*(*k_i_* > 0|*ν_i_* > 0) = 0 for *ν_i_* = {0, *ν*_max_}. This state is denoted as 1 and from now on we have *k_i_* ∈ {0,1}. Thus *P*(*k_i_* = 1|*ν_i_* = *ν*_max_) = 1 − *P*(*k_i_* = 0|*ν_i_* = *ν*_max_) = 1 − *e*^−*R*^. The mutual information can then be calculated as

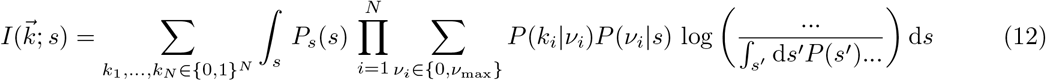

with

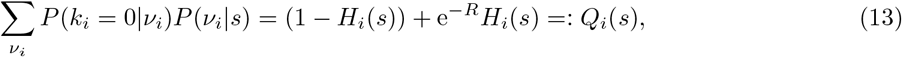

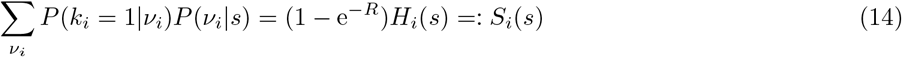

where output noise is denoted as *R*:= *ν*_max_Δ*T*. Taken together, the mutual information for the independent coding channel is

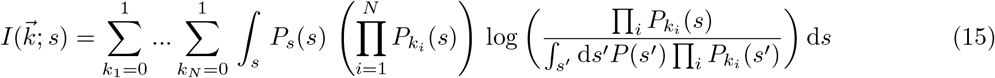

with

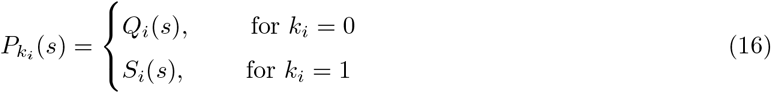

### Lumped-coding channel

Next we turn to the calculation for the input-output kernel *P*(*k|s*) in the case of the lumped-coding channel. For the case of only input input noise where *Q_i_*(*s*) = 1 − *H_i_*(*s*) and *S_i_*(*s*) = *H_i_*(*s*), McDonnell et al. [29,75] explained how *P*(*k|s*) can be calculated using a recursive formula. We extended these calculations to additional Poisson output noise. We write *P*(*k|s*) as *P*(*k|N, s*) and use the notation by McDonnell et al. [29, 75], for which

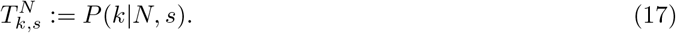

Furthermore, *P_k_i_|s,i_* is defined as the probability of cell *i* firing *k_i_* spikes in a coding window Δ*T* when the stimulus is *s*. With that, one can express the probability of having *k* spikes with *N* cells as the probability of having *k_N_* spikes by the *N*-th neuron multiplied by the probability of having *k* − *k_N_* spikes by the other neurons and taking into account all possibilities of *k_N_* by summing over *k_N_*:

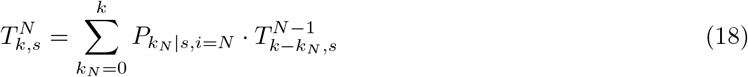

where

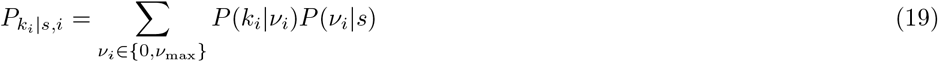

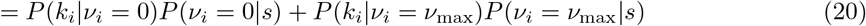

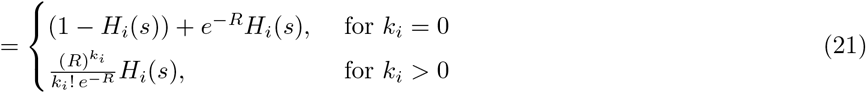

is the probability of cell *i* emitting *k_i_* spikes given stimulus *s*, and

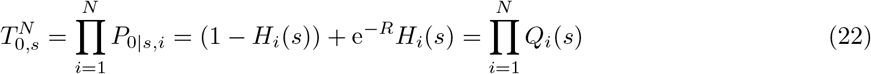

being the probability of having zero spikes with *N* cells, as well as

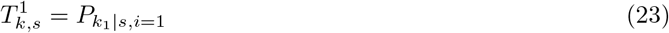

being the probability of having *k* spikes with *N* = 1. Thus for every *k* = 0, 1, 2,… until an upper bound which is determined by the precision one wants to reach, 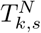 is calculated for every *k_N_* = 0,1,…*k* by using the recursive formula in Eq. 18. This is computationally very expensive and thus we studied only populations with up to *N* = 3 neurons and expected maximum spike count of *R* = 10 (note that calculating just *one P*(*k|s*) for *N* = 3 and *R* =10 requires on the order of 50 000 evaluations of Eq. 19). As with the independent-coding channel, input noise *σ* is included in *H_i_*(*s*) (see Eq. 9) and the output noise level is denoted by *R*.

Our goal is to find the optimal thresholds 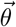 which maximize mutual information for given levels of input and output noise *σ* and *R*:

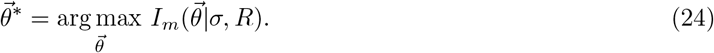

### Optimization procedure

For the calculation of Eq. 4 we performed the integration numerically for both the independent- and the lumped-coding channel. For the independent-coding channel, this numerical integration is the computationally most expensive part of calculating the mutual information. We tested several numerical integration algorithms (Riemann, trapezoid, Romberg, Simpson, and adaptive algorithms) which all lead to very similar results. We performed numerical optimizations using the Nelder-Mead simplex algorithm implemented in the Scipy package [76]. It is a local optimizer which does not rely on estimating the gradient. Gradient based optimizers like the Broyden-Fletcher-Goldfarb-Shanno (BFGS) algorithm rely on calculating or estimating the inverse of the Hessian matrix. For the independent-coding channel this becomes problematic around critical noise values where one eigenvalue of the Hessian approaches zero and thus leads to large numerical imprecisions when inverting the Hessian. For the lumped-coding channel this is unproblematic and for speed purposes we also used an adaptation of the BFGS algorithm [77] implemented in Scipy. In order to spot possible local maxima – which are especially prevalent for large *N* – we applied a grid of initial conditions. After some trials it was possible to estimate what form of initial conditions lead to local maxima. Additionally, potential local maxima could in general be easily spotted and checked.

The heavy numerical procedure limited our analysis to small population sizes with a maximum of three neurons in the case of the lumped-coding channel, and six neurons in the case of the independent-coding channel.

### Generalized normal distribution

The generalized normal distribution (GND) is given by [41]

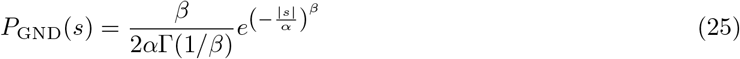

where Γ(*z*) is the gamma function given by

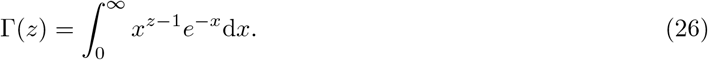

The parameter *β* determines the kurtosis, particular values being *β* = 1 (for the Laplace distribution), *β* = 2 (the standard normal distribution) and *β* → ∞ (the uniform distribution). The variance of the GND is

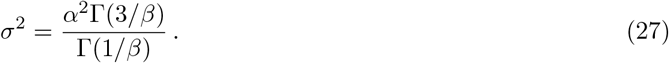

The effective tuning curve of Eq. 9 is in this case given as

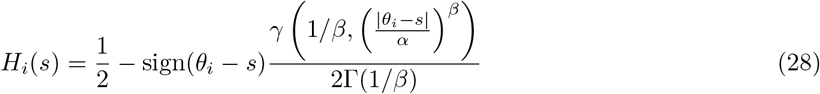

where *γ*(*x, y*) is the lower incomplete gamma function defined as

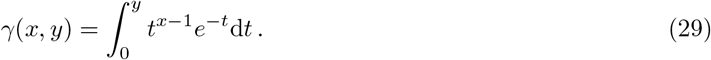

### Local curvature of information landscape

To investigate the curvature of the information landscape, we numerically calculated the Hessian matrix of the mutual information (using the Python package Numdifftools) at optimal thresholds and performed eigendecomposition. The Hessian matrix is defined as

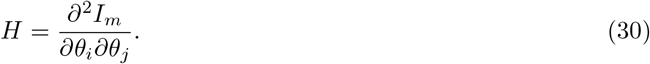

Its eigenvectors give the directions of principal curvatures and the respective eigenvalues quantify the curvature in these directions.

### Data fitting

We modeled ANF tuning curves as binary neurons, each neuron i with threshold *θ_i_* so that if the stimulus (here, sound intensity at a given frequency) is higher (lower) than *θ_i_*, the firing rate is *r_m,i_*(*r_i_*). Here *r_i_* denotes the spontaneous firing rate (SR) of the neuron, and *r_m,i_* denotes its maximal firing rate. The addition of Gaussian input noise with mean 0 and standard deviation *σ_i_* transforms the effective tuning curve of the neuron into a sigmoid given by the equation (Fig. 1A & Fig. 7A):

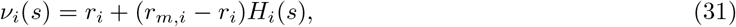

where *H_i_* denotes the complementary error function (Eq. 11). To analyze the experimentally recorded ANF tuning curves from ref. [8], we first fit all the tuning curves with Eq. 31. We used the approach from Balasooriya et al. [78] to identify a single outlier in the distribution of normalized SR, *r_i_*/*r_m,i_*. Upon removing the outlier, we pooled the measured tuning curves from the same ANF, and fitted each with our sigmoidal function (Fig. 7A). This resulted in 148 tuning curves from 24 animals. To divide the tuning curves into two classes, since the distributions of normalized SR and the dynamic range are not center-symmetric, we calculated the cumulative distribution functions, *F*(*r_i_*/*r_m,i_*) and *F*(*σ_i_*) (Fig. S4).

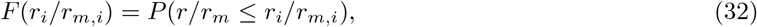

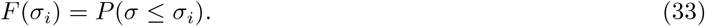

When *F*(*r_i_*/*r_m,i_*) > *F*(*σ_i_*), neuron *i* was classified as ‘Type 1,’ characterized by a higher SR, smaller dynamic range and a lower firing threshold. Otherwise neuron i was classified as ‘Type 2,’ characterized by a lower SR, broader dynamic range and a higher firing threshold (e.g. Fig. 7B).

After fitting, we extracted the average values of all parameters for each type. The ‘Type 1’ neuron had a high normalized SR (*r*_1_/*r_m_* = 0.16) and a dynamic range of *σ*_1_ = 0.34; while the ‘Type 2’ neuron had a very low normalized SR (*r*_2_/*r_m_* = 0.036, which we approximated to 0) and a dynamic range of *σ*_2_ = 0.53. These dynamic ranges were normalized to the stimulus distribution of natural sound frequencies used in the optimization, which is assumed to be a Gaussian distribution with mean 30 dB and standard deviation of 12.5 dB [79]. The maximum expected spike count *R* – the product of average maximal firing rate of the ANFs and the coding time window, Δ*T* = 50 ms – was 13.8.

## Acknowledgements

We thank M. Charles Liberman (MIT) for providing us with the data set of auditory nerve fibers recorded in the mouse. We also thank the entire ‘Computation in Neural Circuits’ group for helpful discussions, and Marcel Jungling and Sebastian Onasch for feedback on the manuscript.

## Supplementary Figures

**Figure S1.**
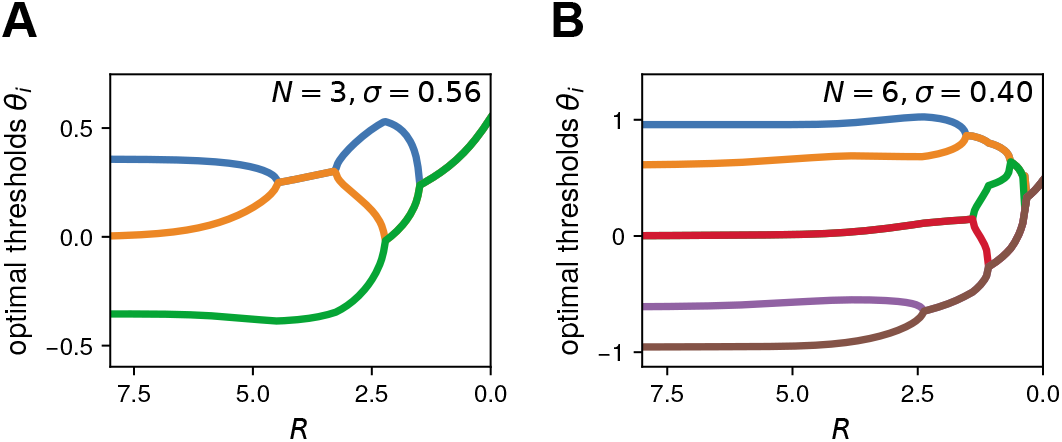
Non-monotonicity of the number of distinct optimal thresholds with output noise level *R*. **A.** For *N* = 3 neurons there is a non-monotonicity of the number of distinct optimal thresholds with output noise for a relatively small input noise parameter range (0.54 < *σ* < 0.6, see Fig. 4A). For high output noise (low R), first the upper two thresholds merge, before they split again with decreasing output noise and for even lower output noise the middle threshold merges with the lower threshold. **B.** For *N* = 6 neurons a similar transformation of thresholds happens, where the two middle thresholds split with decreasing output noise, thus increasing number of distinct optimal threshold.

**Figure S2.**
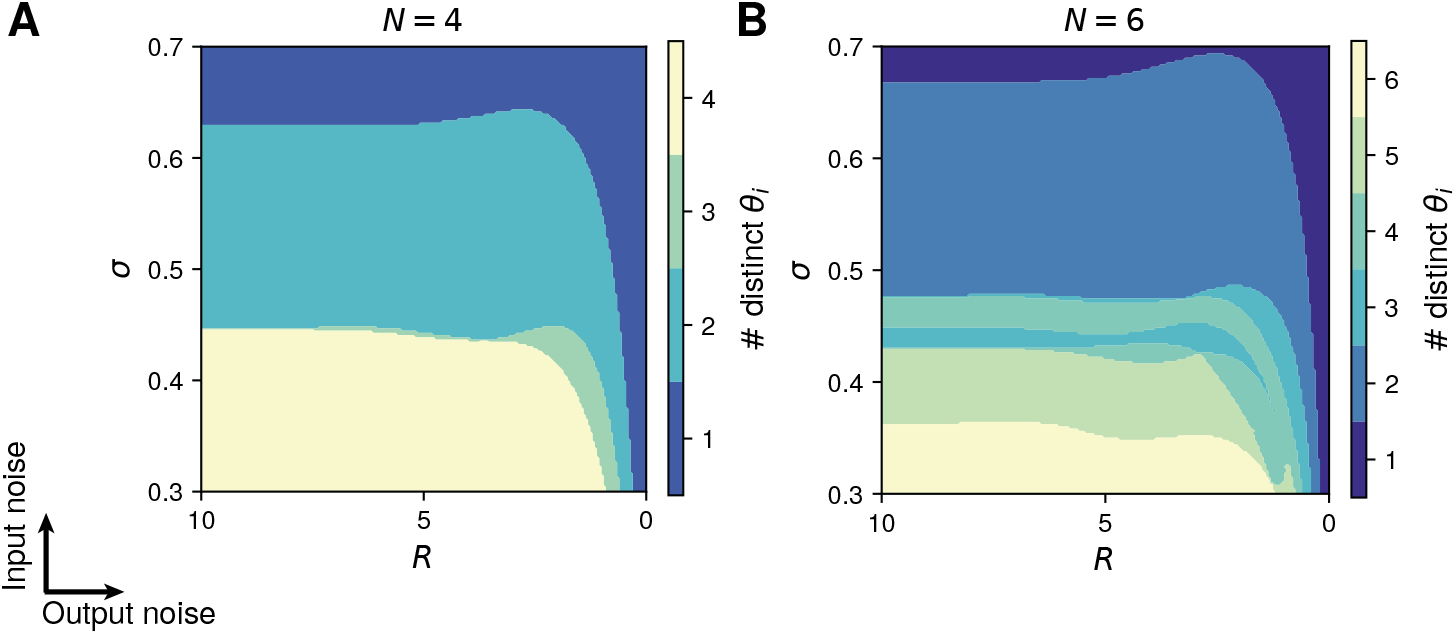
Optimal thresholds for higher number of neurons in the case of the independent-coding channel. **A.** Number of distinct optimal thresholds for *N* = 4 cells depending on input noise *σ* and output noise *R*. **B.** As in A but for *N* = 6.

**Figure S3.**
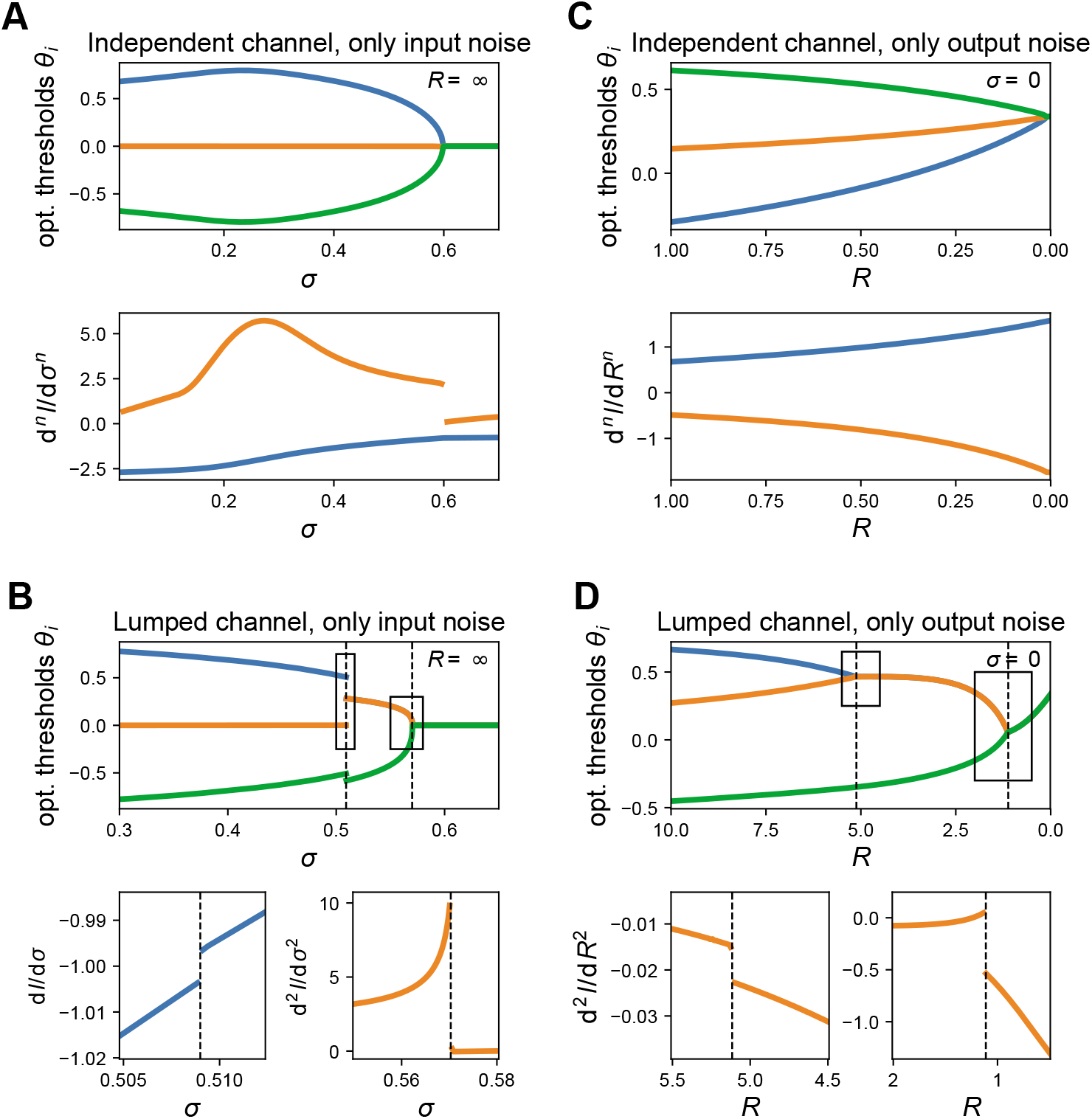
Threshold differences as phase transitions in the limit of just one noise source. **A**. Threshold bifurcations for the independent-coding channel with respect to input noise *σ* for vanishing output noise. The derivatives of mutual information with respect to input noise indicate a second-order phase transition. **B**. As in A but for lumped-coding channel. There is first-order phase transition for low noise (left inset) and a second-order phase transition for high noise (right inset). **C**. Independent-coding channel with vanishing input noise. No phase transition is visible since the “bifurcation” happens in the limit of infinite output noise. **D**. Lumped-coding channel with vanishing input noise exhibits second-order phase transitions.

**Figure S4.**
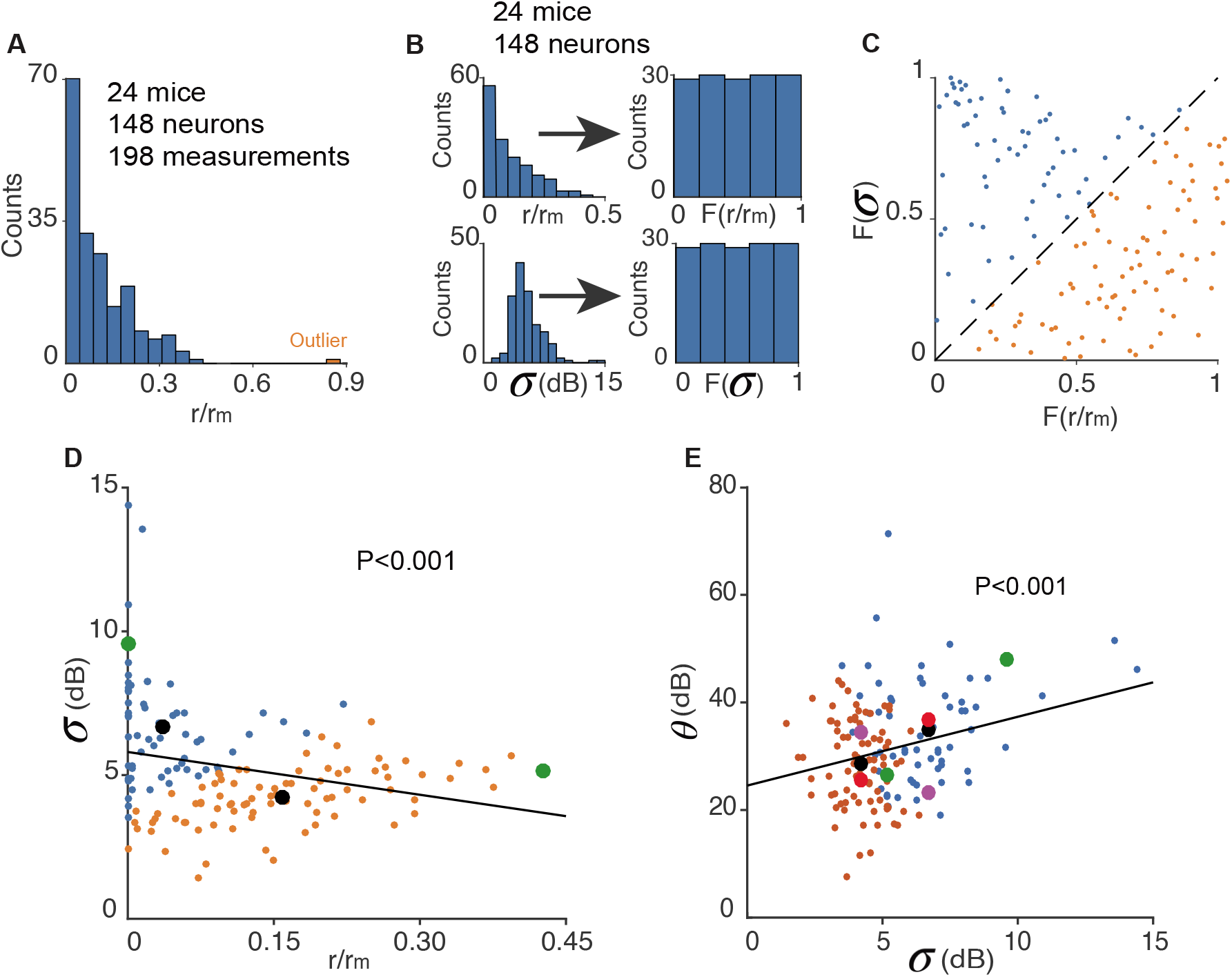
Analysis of auditory nerve tuning curves of mice. **A.** Distribution of normalized SR. An outlier is identified and marked in orange. **B.** Transforming normalized SR (*r/r_m_*) and dynamic range *σ* into cumulative distribution functions. **C.** A diagram showing the method to classify neurons into two types. **D.** Scatter plot and linear fit between normalized SR and dynamic range *σ*. Black dots denote the ‘center of mass’ within each ‘type’, and the green dots show the values of example neurons in Fig. 7B. **E.** As in D but for the relationship between *σ* and threshold *θ*. Magenta dots and red dots denote where mutual information is maximized, with corresponding *σ*_1_ and *σ*_2_ as the black dots. Average firing rate corresponding to red dots are lower.

**Figure S5.**
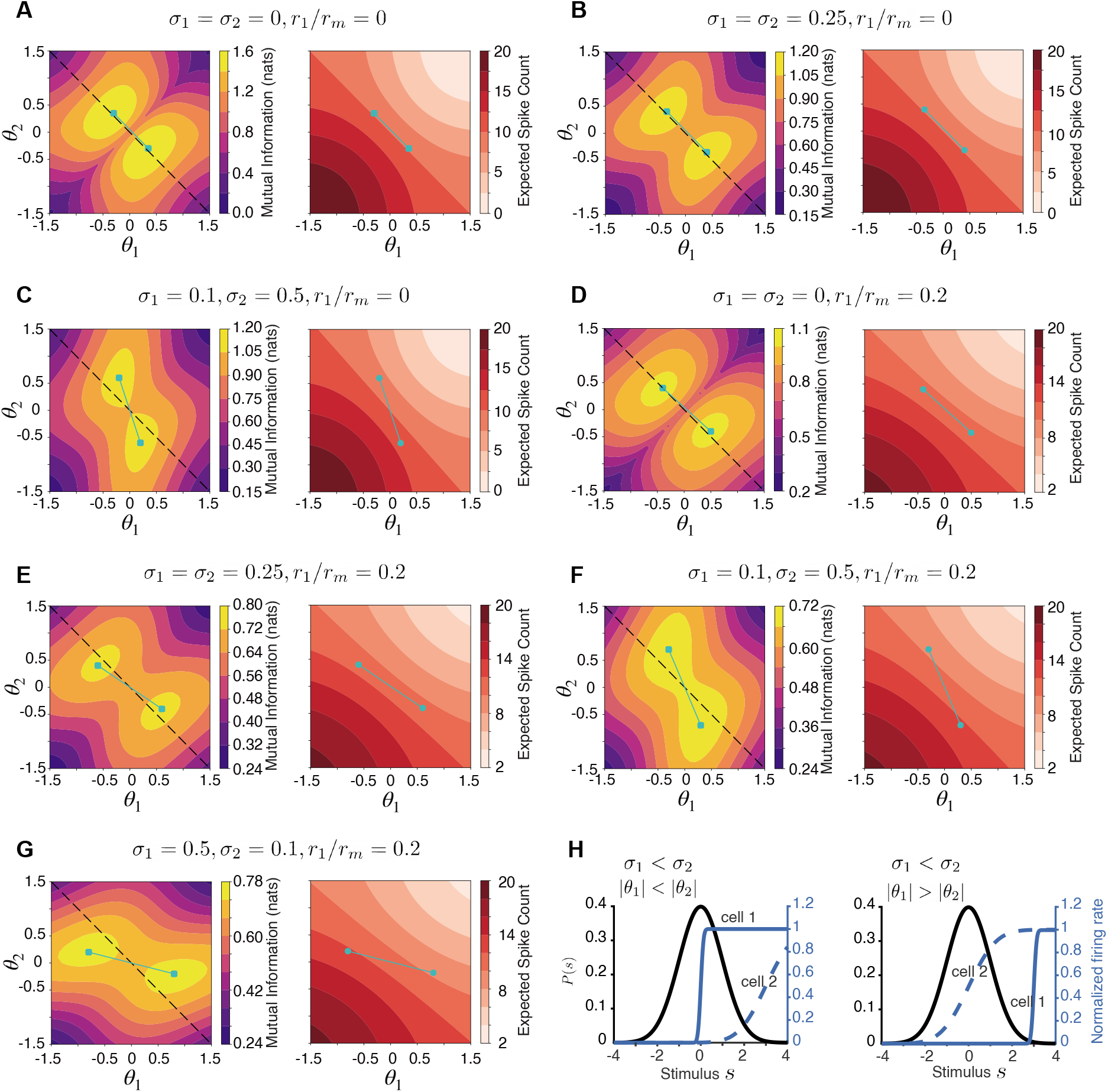
Contour plots of mutual information and average firing rate, with different combination of *σ*_1_, *σ*_2_ and *r*_1_/*r_m_*. In each panel, the left plot corresponds to mutual information and the right one shows average firing rate. Cyan dots show optimal thresholds (*θ*_1_, *θ*_2_) which maximize mutual information. Maximal spike count is set to *R* =10 for every panel. **A.** Both neurons have zero input noise *σ*_1_ = *σ*_2_ = 0, and zero spontaneous rate *r*_1_ /*r_m_* = 0. **B.** The two neurons have identical but nonzero input noise *σ*_1_ = *σ*_2_ = 0.25, and zero spontaneous rate *r*_1_/*r_m_* = 0. **C.** The two neurons have two different and nonzero input noise *σ*_1_ = 0.1, *σ*_2_ = 0.5, and zero spontaneous rate *r*_1_/*r_m_* = 0. **D.** Both neurons have zero input noise *σ*_1_ = *σ*_2_ = 0, and nonzero spontaneous rate *r*_1_/*r_m_* = 0.2. **E.** The two neurons have identical but nonzero input noise *σ*_1_ = *σ*_2_ = 0.25, and non-zero spontaneous rate *r*_1_/*r_m_* = 0.2. **F.** The two neurons have two different and nonzero input noise *σ*_1_ = 0.1, *σ*_2_ = 0.5, and nonzero spontaneous rate *r*_1_/*r_m_* = 0.2. **G.** *σ*_1_ = 0.5, *σ*_2_ = 0.1, *r*_1_/*r_m_* = 0.2. **H.** The mechanism behind symmetry breaking of the mutual information landscape. The case (left) where the neuron with the larger input noise has a larger threshold located in the region where the stimulus rarely occurs is more efficient than in the case (right) where the neuron with the larger input noise has a smaller threshold near the stimulus mean where its dynamic range covers a large range of possible stimuli.

**Figure S6.**
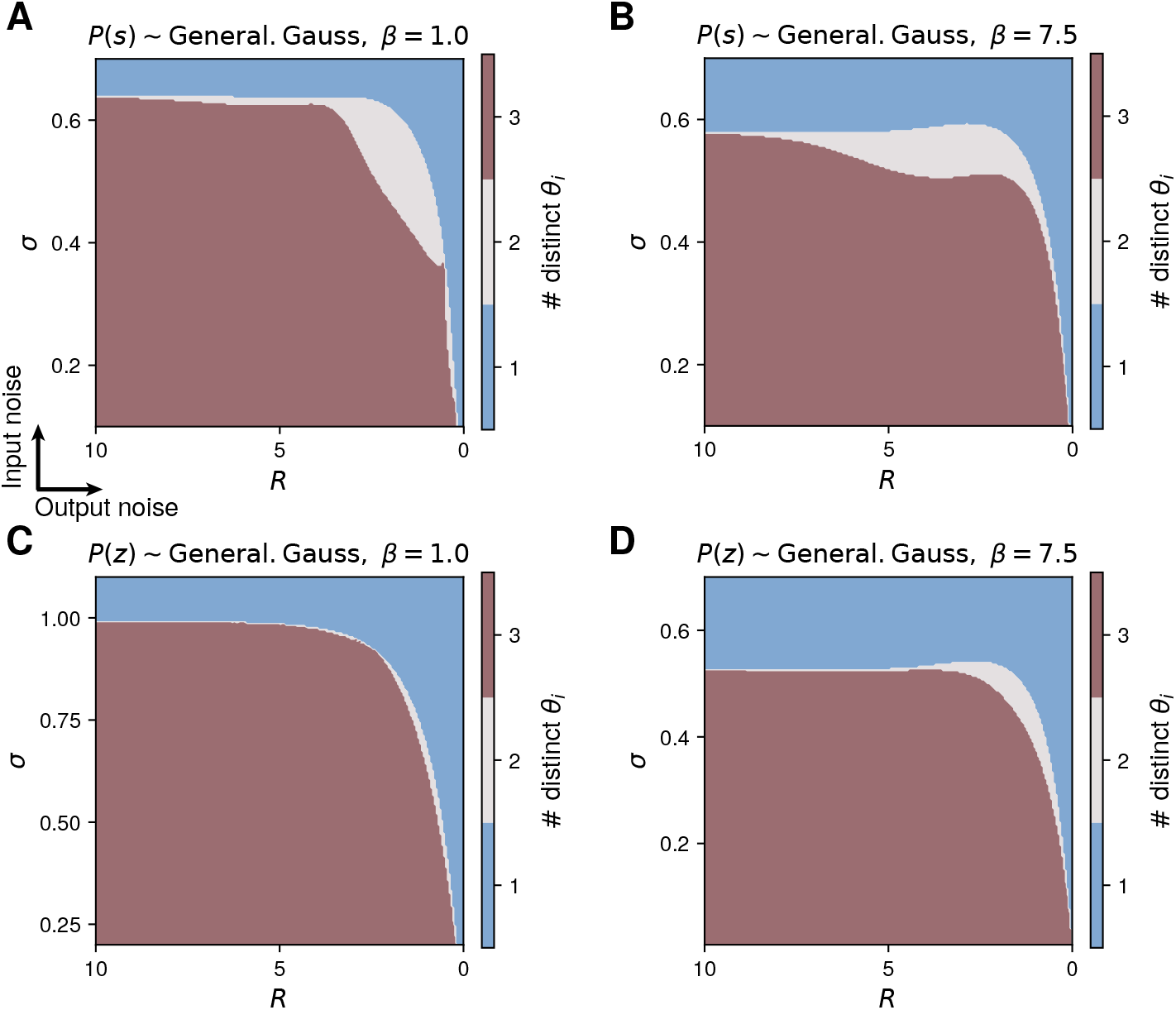
Number of distinct optimal thresholds when using input and noise distributions different from Gaussian. Instead of a Gaussian stimulus and noise distribution we also used a generalized normal distribution and varied the kurtosis. **A.** Laplacian (having high kurtosis) as input distribution. **B.** Input distribution with low kurtosis (similar to uniform). **C.** Laplace distribution as noise distribution. **D.** Noise distribution with low kurtosis.

